## Supplementary material for "CRISPR-Cas12a exploits R-loop asymmetry to form double-strand breaks": Supp_Figures_2-8.pdf

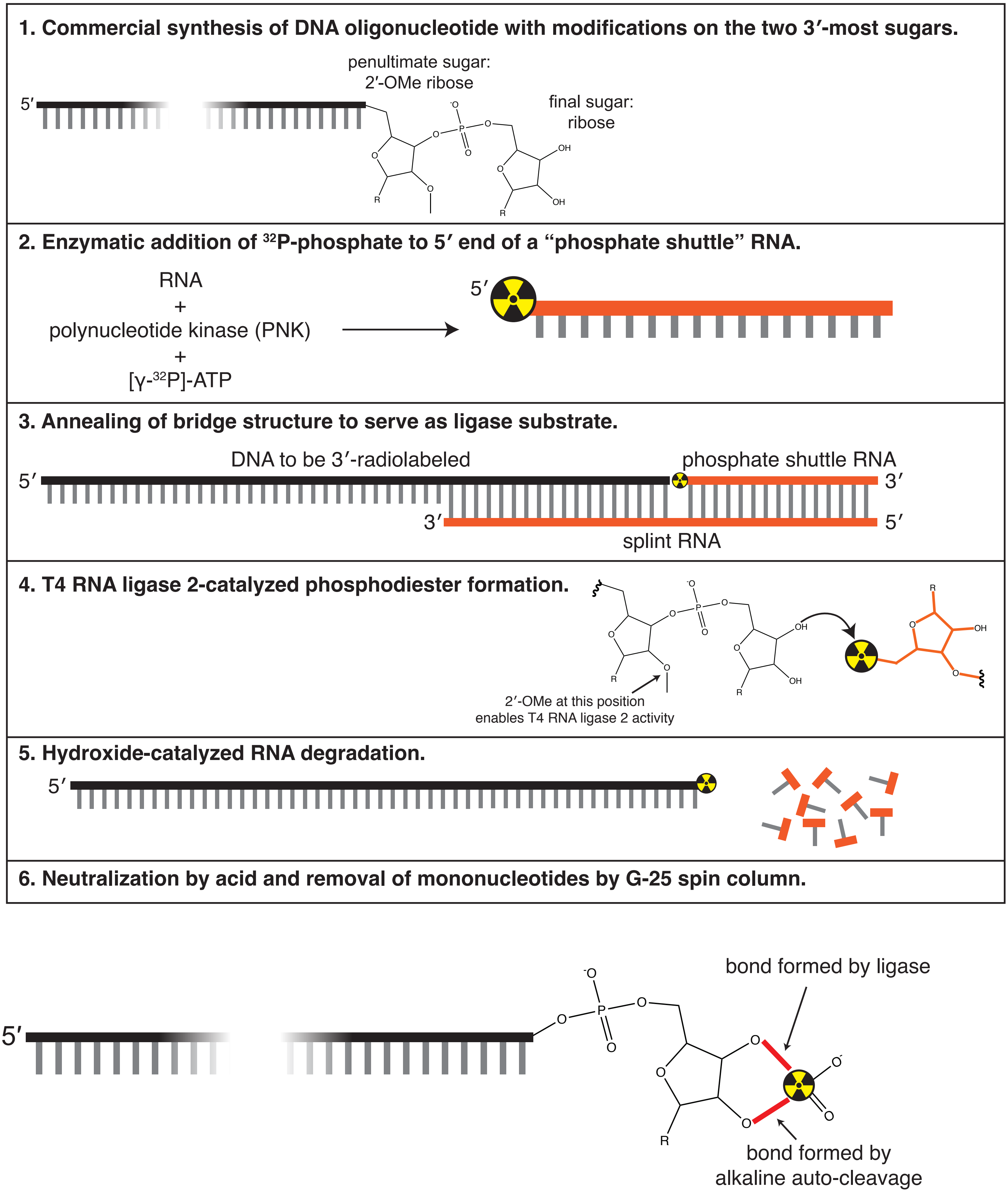


**Supplementary Figure 2A.**

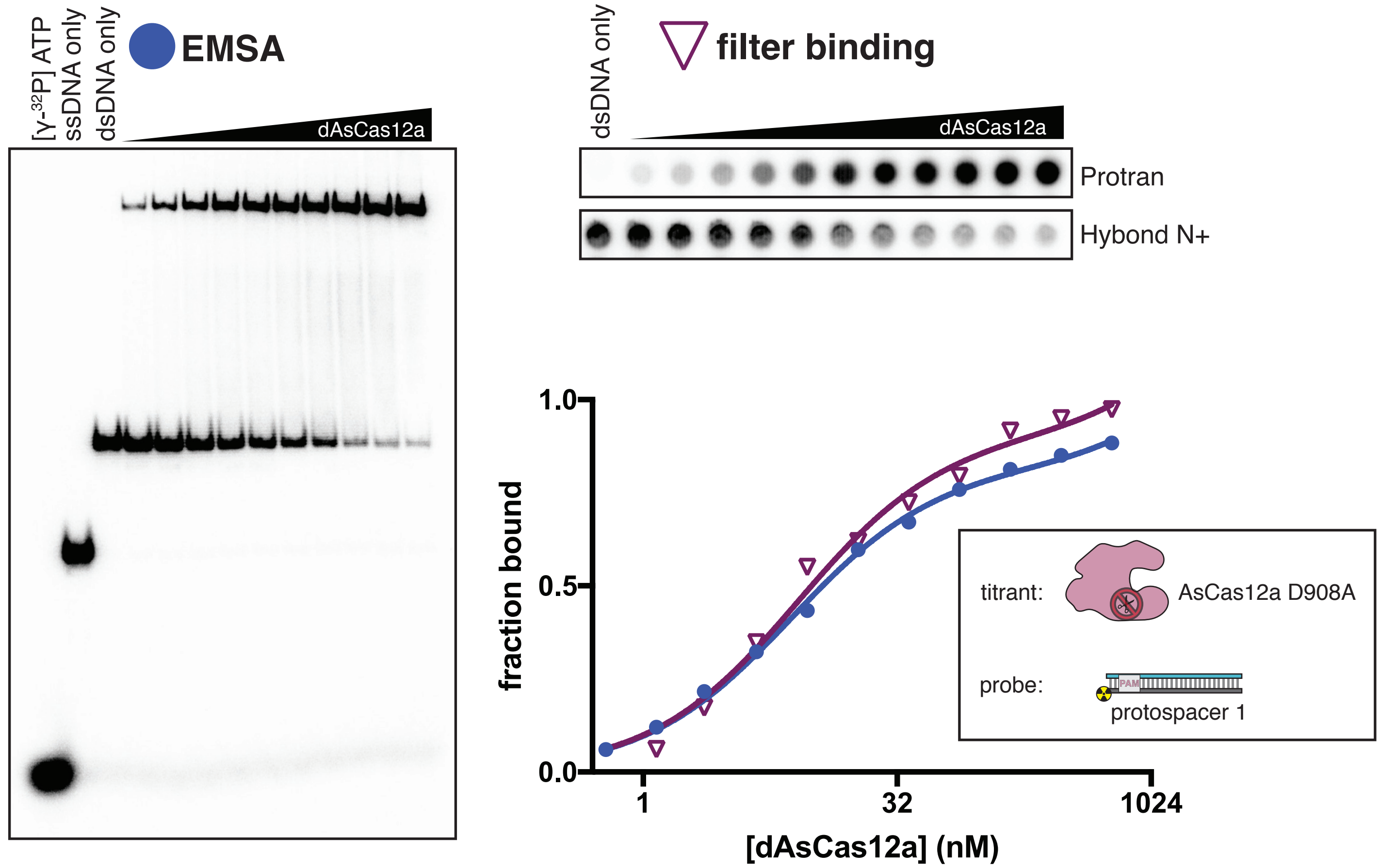


**protospacer 1:** PAM | complementary to crRNA spacer | R-loop flank (RLF)

```
TS  3′-CAGTATTACTAAAATAGAAGACCTAACAACATTCGTCGTAAACTCGTTTTTAGACAACG-5′
NTS 5′-GTCATAATGATTTTATCTTCTGGATTGTTGTAAGCAGCATTTGAGCAAAAATCTGTTGC-3′
```

**protospacer 2:** PAM | complementary to crRNA spacer | R-loop flank (RLF)

```
TS  3′-CCGTATCACTGAAATAGAAGACCTAACAACATTCGATATATATTCGTGCTCGGACAACG-5′
NTS 5′-GGCATAGTGACTTTATCTTCTGGATTGTTGTAAGCTATATATAAGCACGAGCCTGTTGC-3′
```

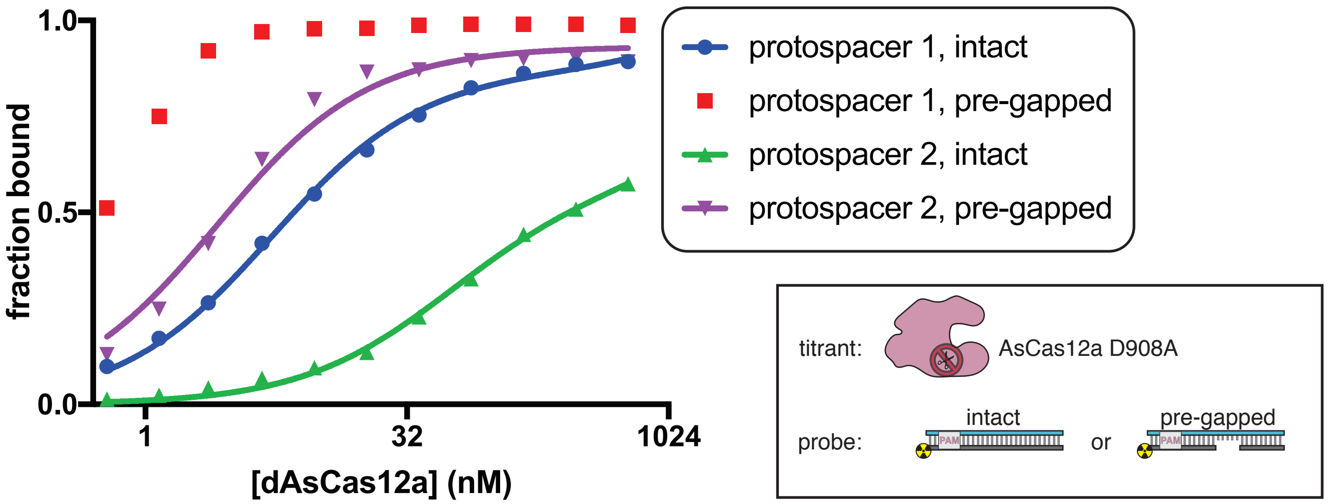


**Supplementary Figure 2B.**

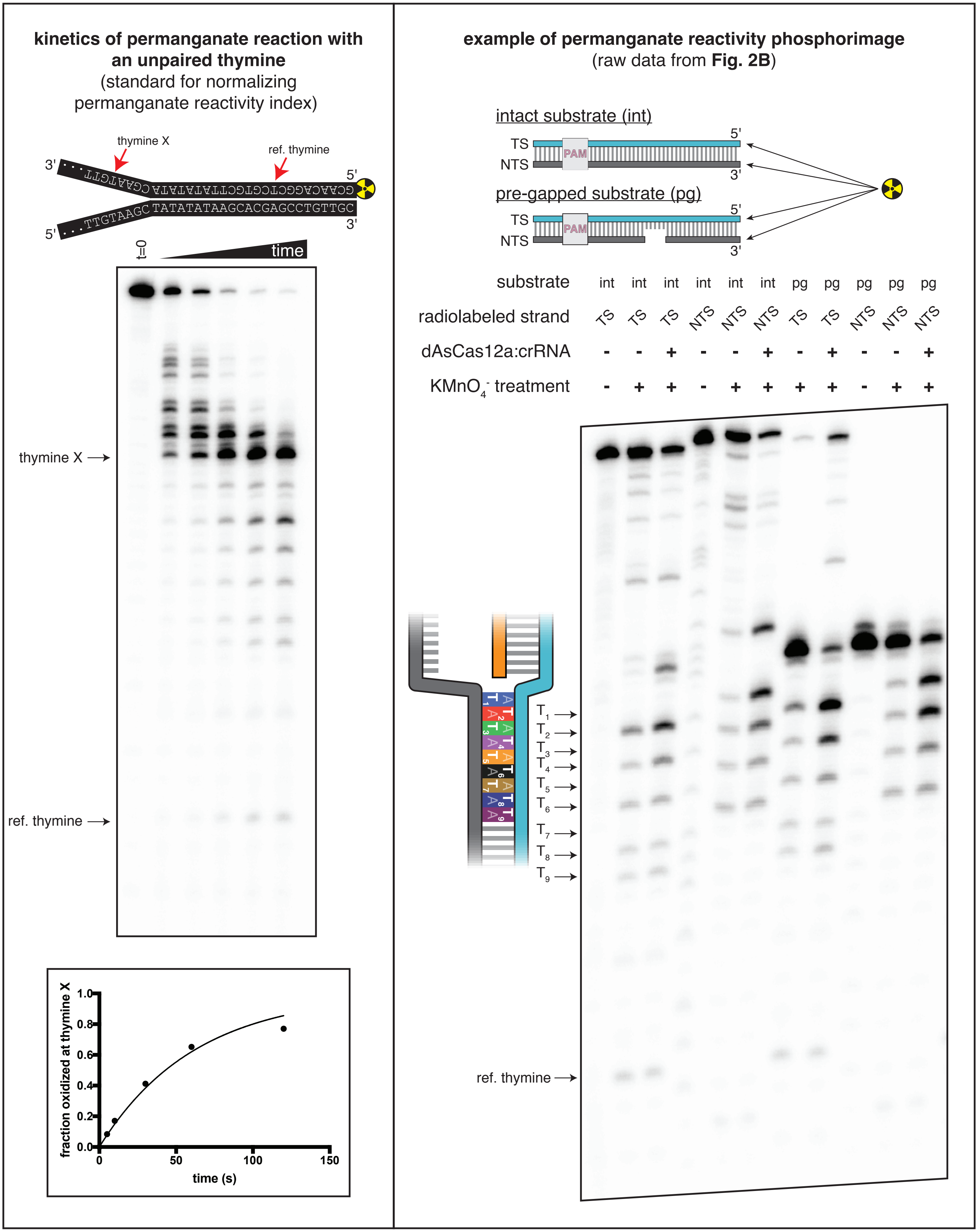


**Supplementary Figure 2C.**

**protospacer 2:** PAM, A/T tract

TS 3′-CCGTATCACTGAAATAGAAGACCTAACAACATTCGATATATATTCGTGCTCGGACAACG-5′

NTS 5′-GGCATAGTGACTTTATCTTCTGGATTGTTGTAAGCTATATATAAGCACGAGCCTGTTGC-3′

**protospacer 3:** PAM, A/T tract

TS 3′-CCGTATCACTGAAATAGAAGACCTAACAACATTCGATATATATTCGTGCTCGGACAACG-5′

NTS 5′-GGCATAGTGACTTTA**AGAAGACCTAACA** GCTATATATAAGCACGAGCCTGTTGC-3′

**spacer of crRNA 3:** 5′-UCUUCUGGAUUGUUGUAAGC-3′

**protospacer 4:** PAM, A/T tract

TS 3′-CCGTATCACTGAAATAGAAGACCTAACAACATTGCATATATATTCGTGCTCGGACAACG-5′

NTS 5′-GGCATAGTGACTTTA**AGAAGACCTAACA** CGTATATATAAGCACGAGCCTGTTGC-3′

**spacer of crRNA 4:** 5′-UCUUCUGGAUUGUUGUAACG-3′

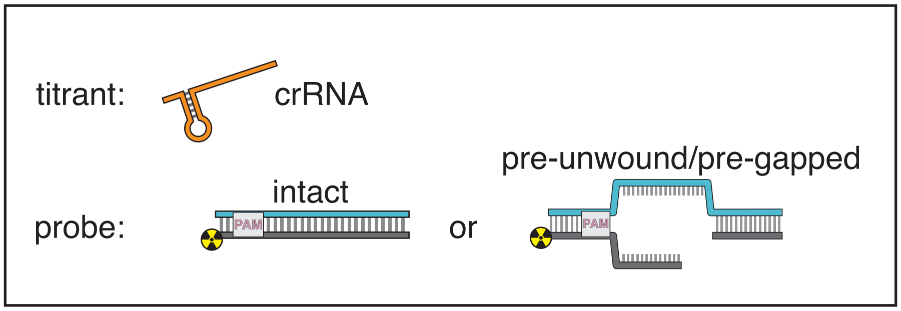


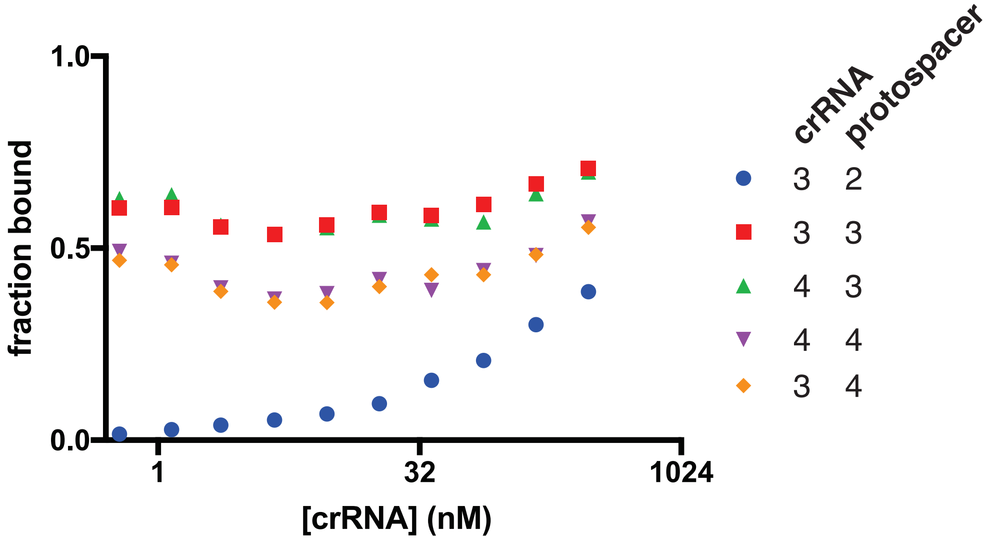


**Supplementary Figure 3A.**

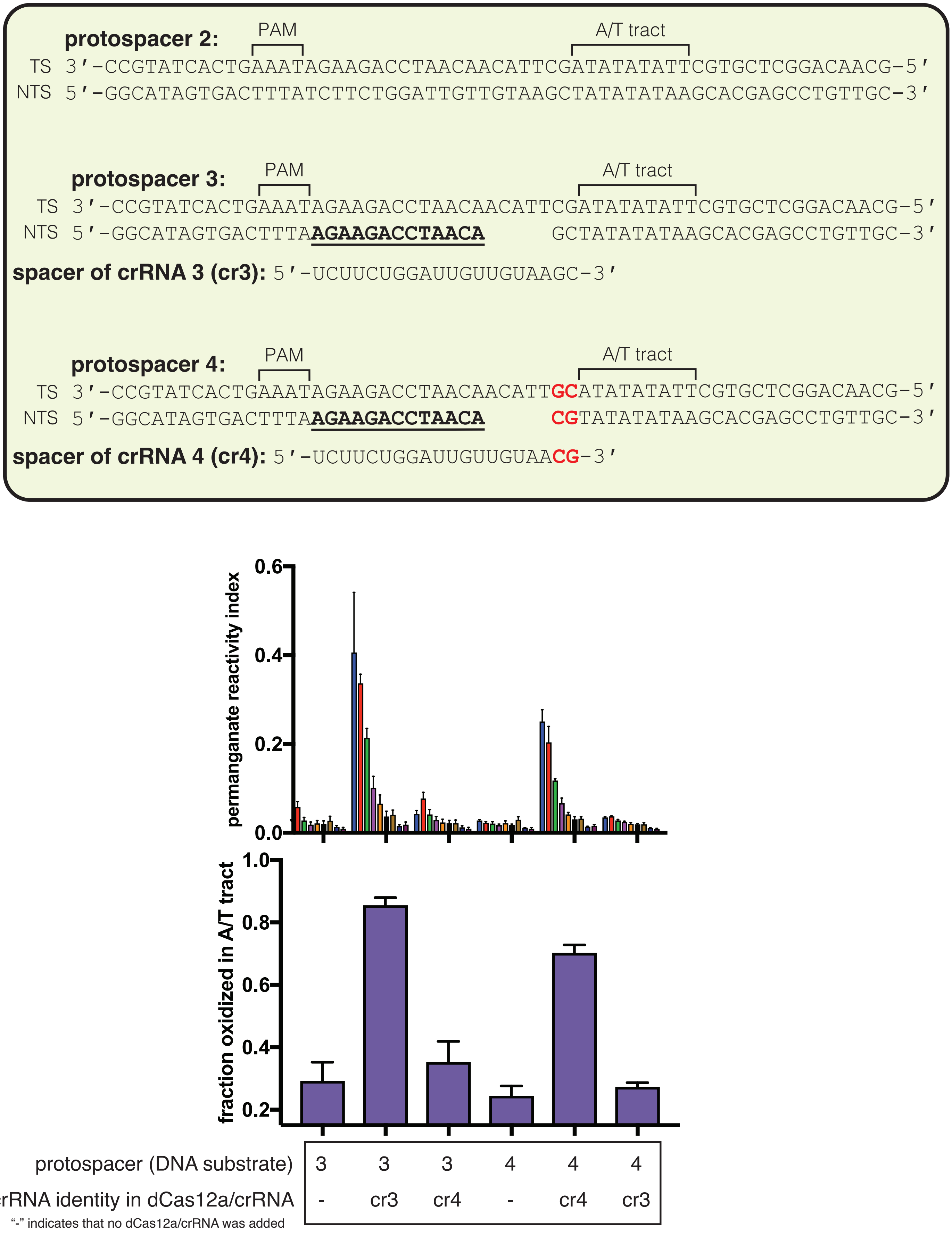


**Supplementary Figure 3B.**

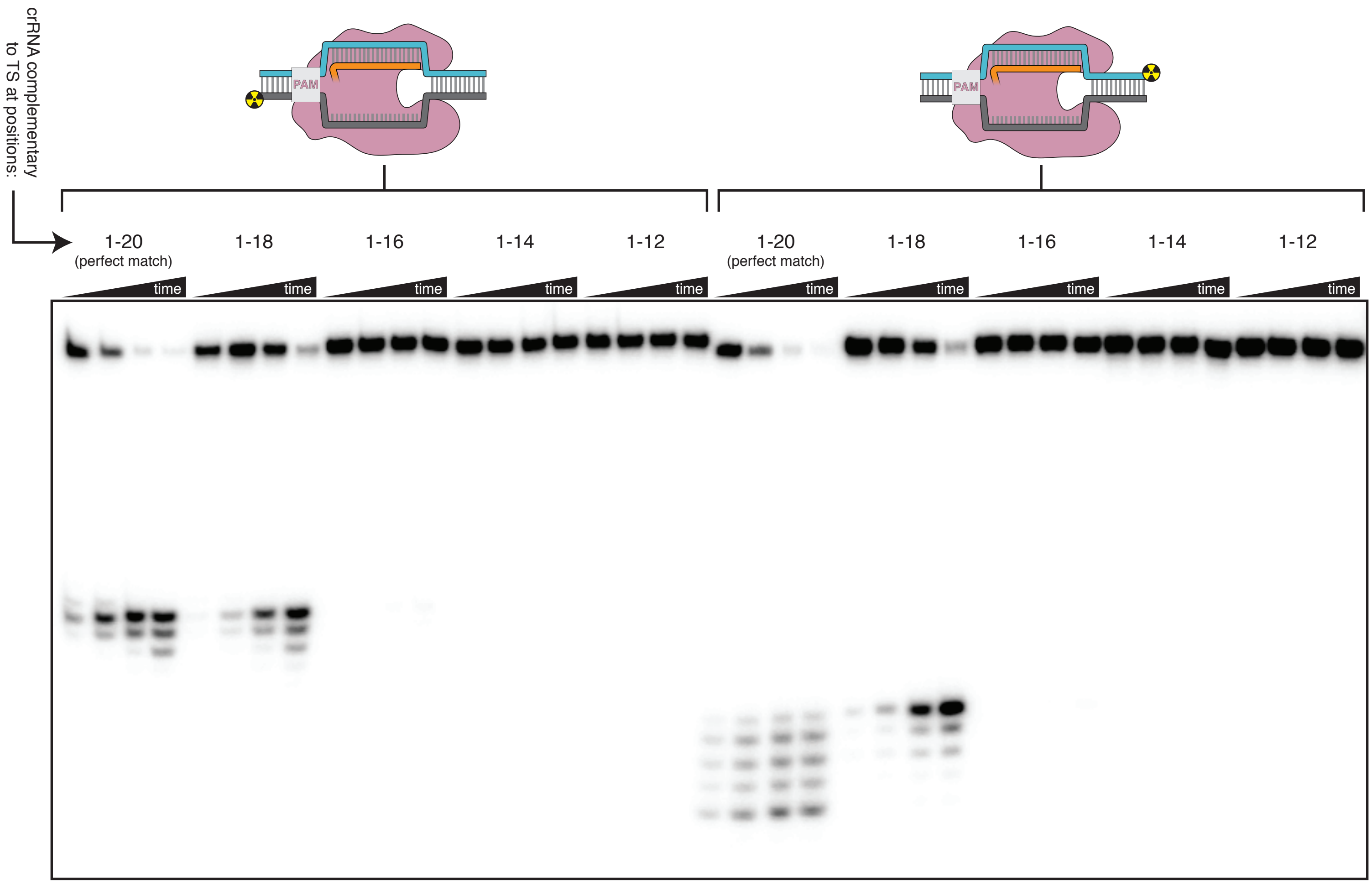


**Supplementary Figure 3C.**

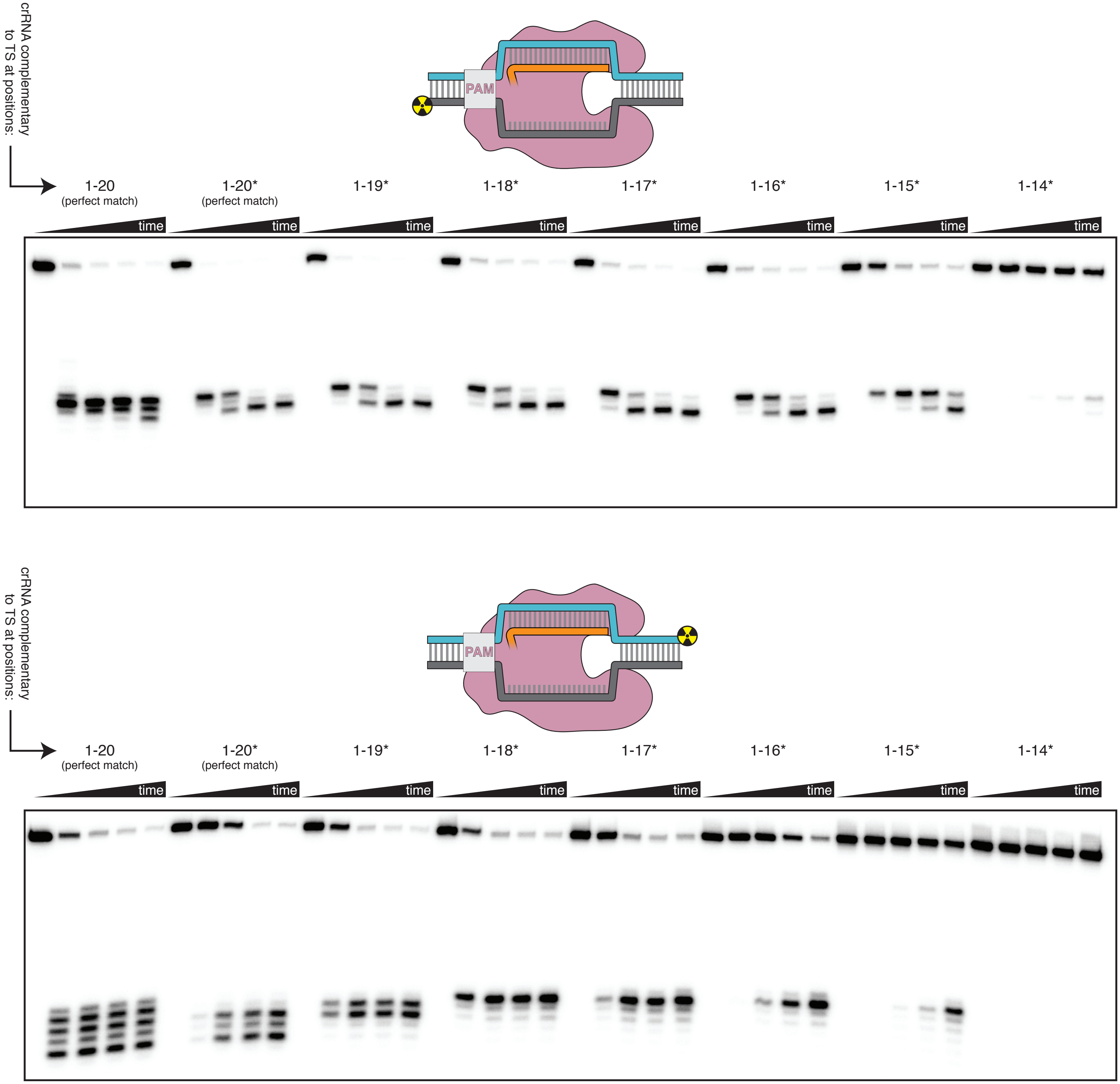


**Supplementary Figure 3D.**

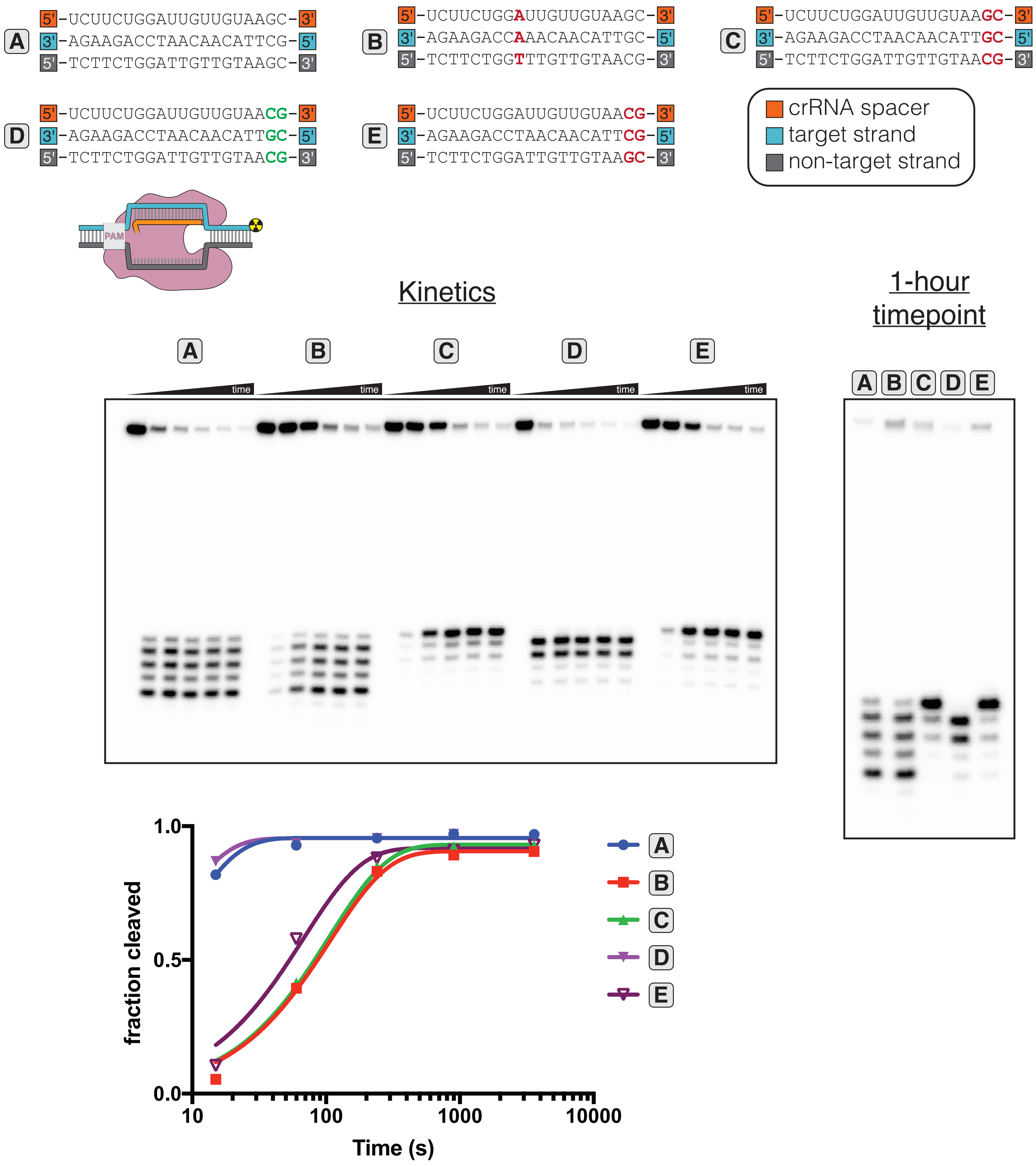


**Supplementary Figure 3E.**

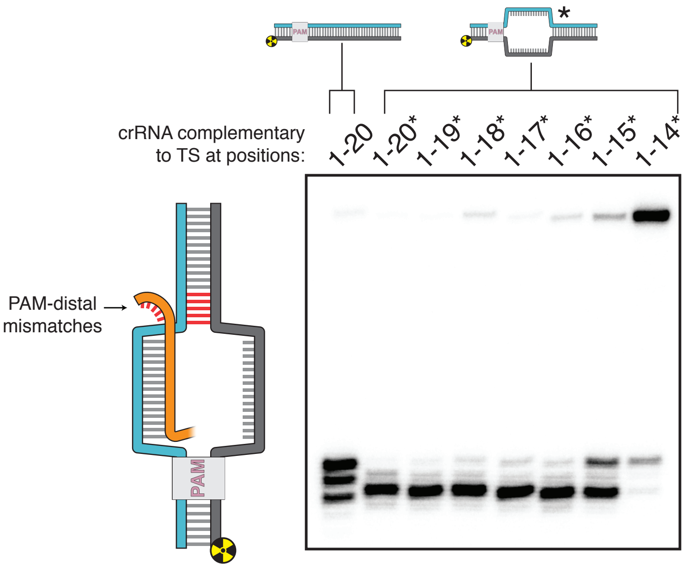


**Supplementary Figure 3F.**

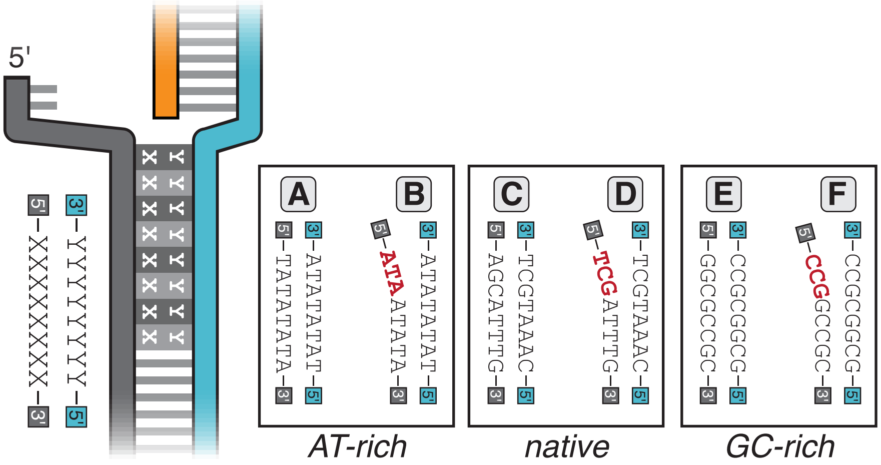


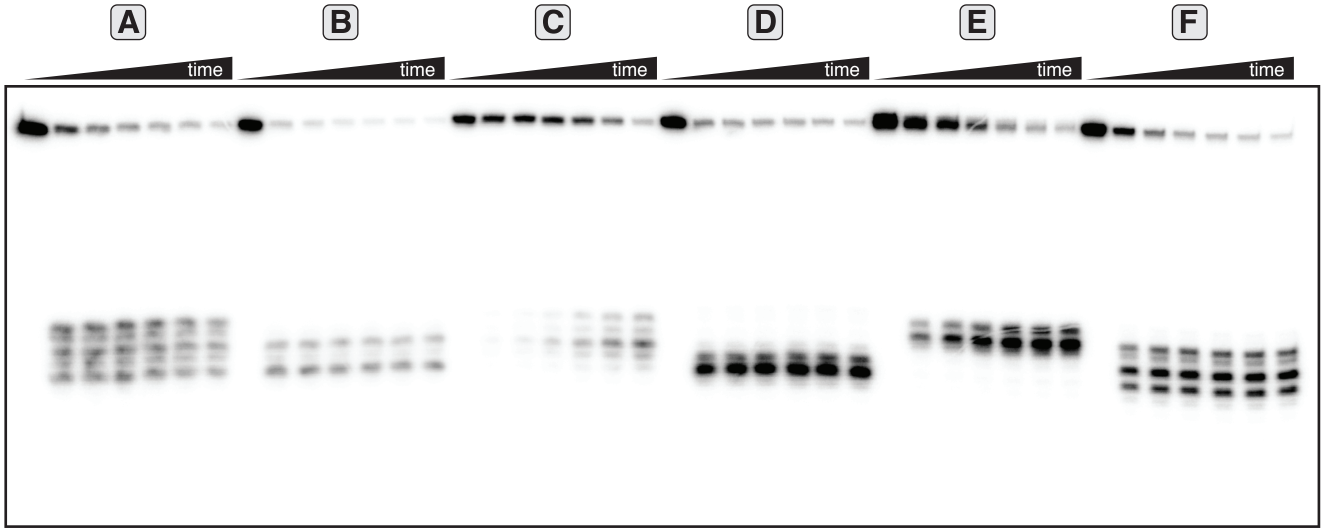


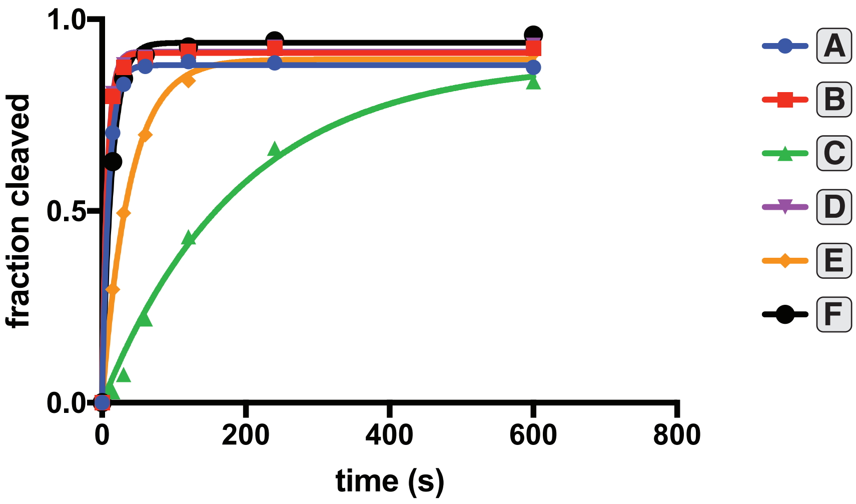


**Supplementary Figure 3G.**

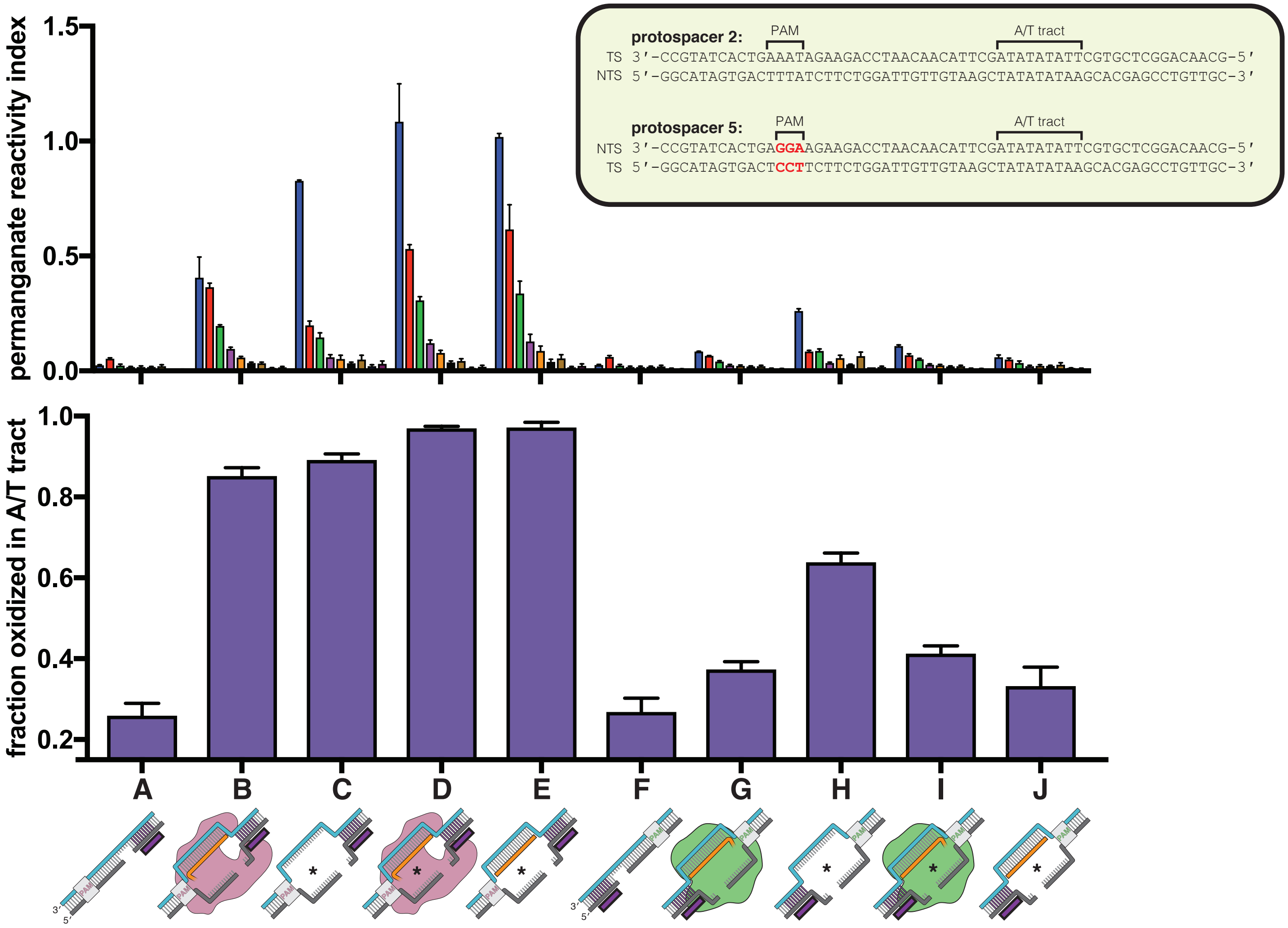


| | DNA construct | additional components |
|---|---|---|
| A | pre-gapped protospacer 2 | - |
| B | pre-gapped protospacer 2 | dAsCas12a + crRNA (20-nt spacer complementary to TS) |
| C | pre-gapped, pre-unwound protospacer 2 | - |
| D | pre-gapped, pre-unwound protospacer 2 | dAsCas12a + crRNA (20-nt spacer complementary to TS) |
| E | pre-gapped, pre-unwound protospacer 2 | 20-nt RNA oligo complementary to TS |
| F | pre-gapped protospacer 5 | - |
| G | pre-gapped protospacer 5 | dSpCas9:sgRNA (20-nt spacer complementary to TS) |
| H | pre-gapped, pre-unwound protospacer 5 | - |
| I | pre-gapped, pre-unwound protospacer 5 | dSpCas9:sgRNA (20-nt spacer complementary to TS) |
| J | pre-gapped, pre-unwound protospacer 5 | 20-nt RNA oligo complementary to TS |

**Supplementary Figure 4A.**

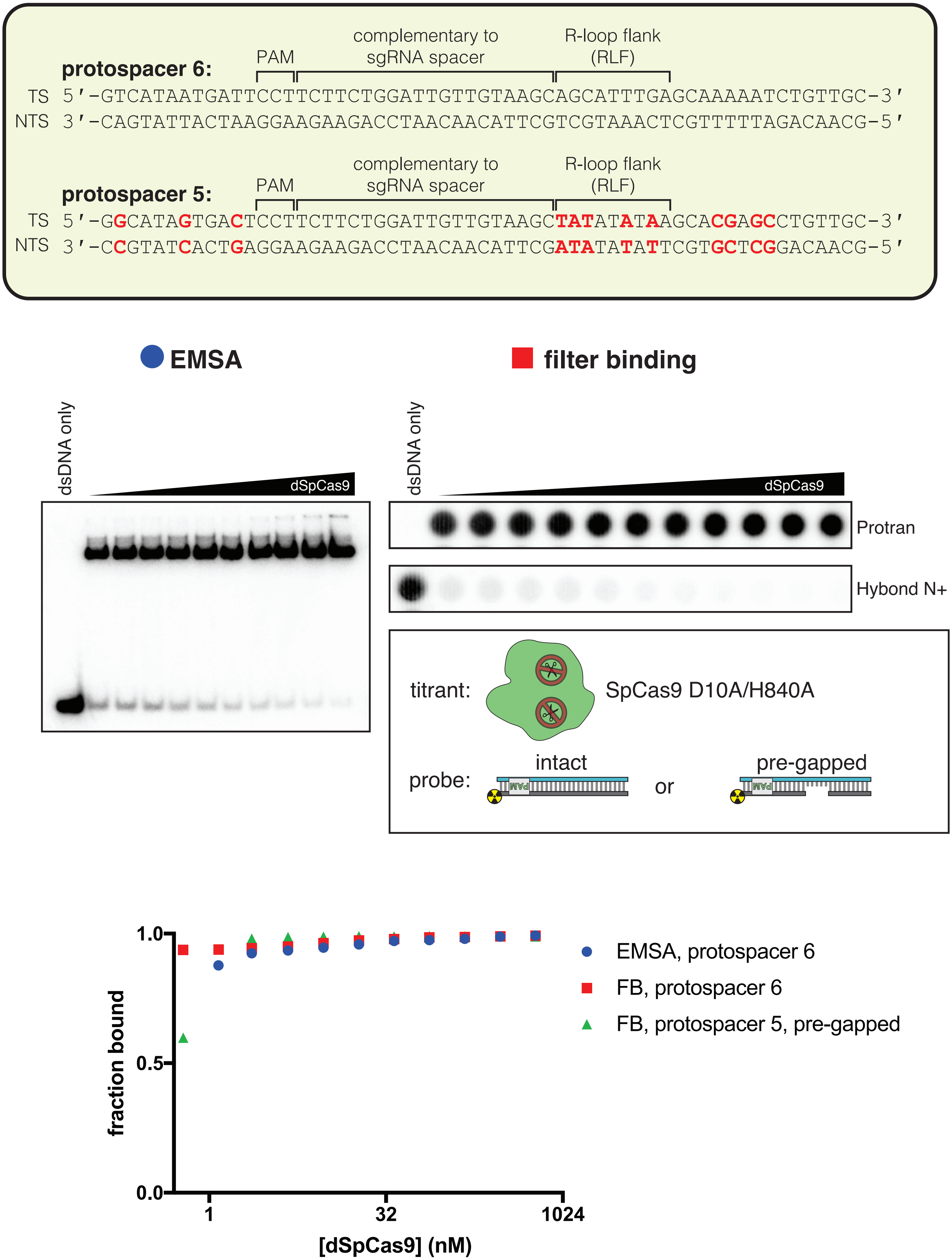


**Supplementary Figure 4B.**

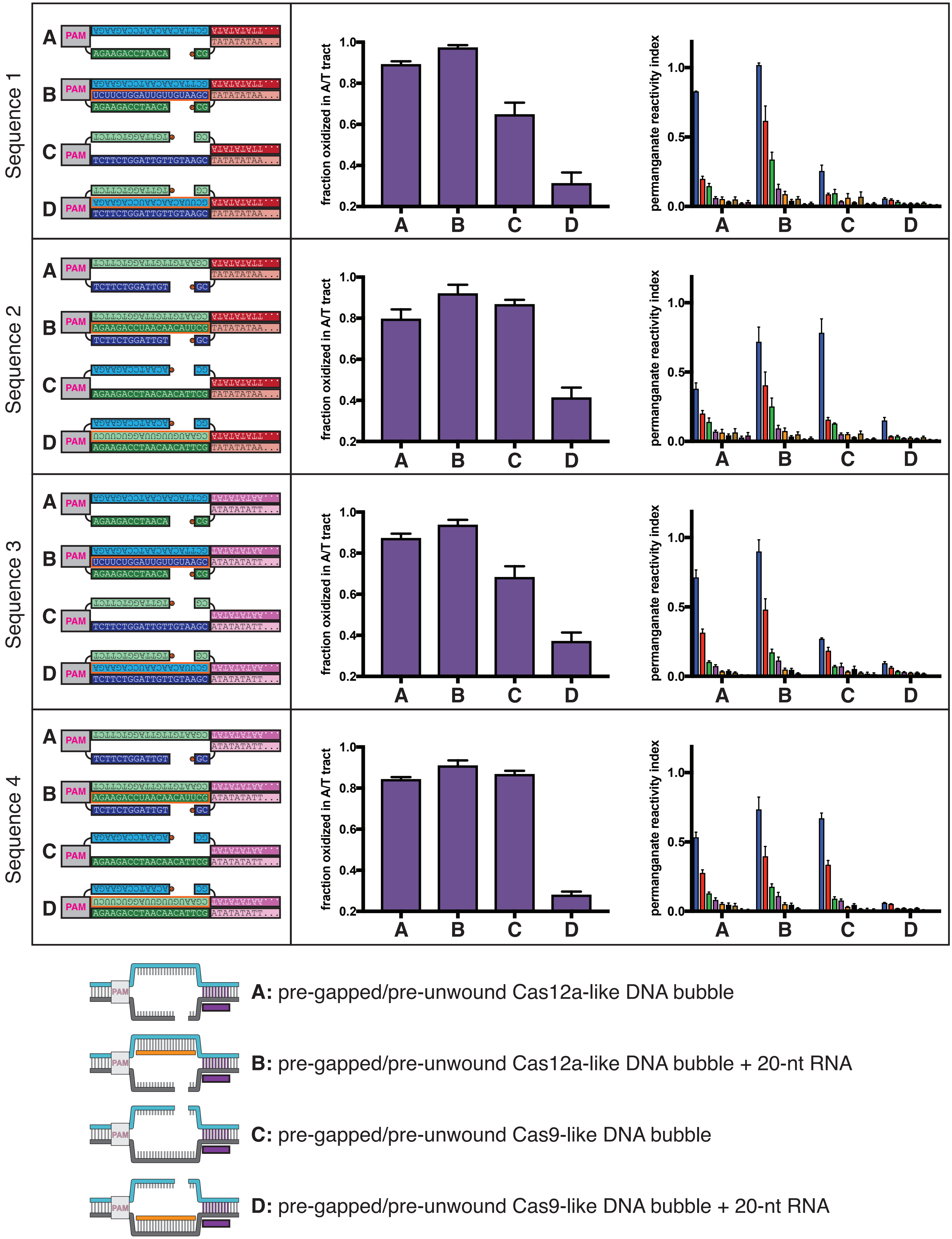


**Supplementary Figure 4C.**

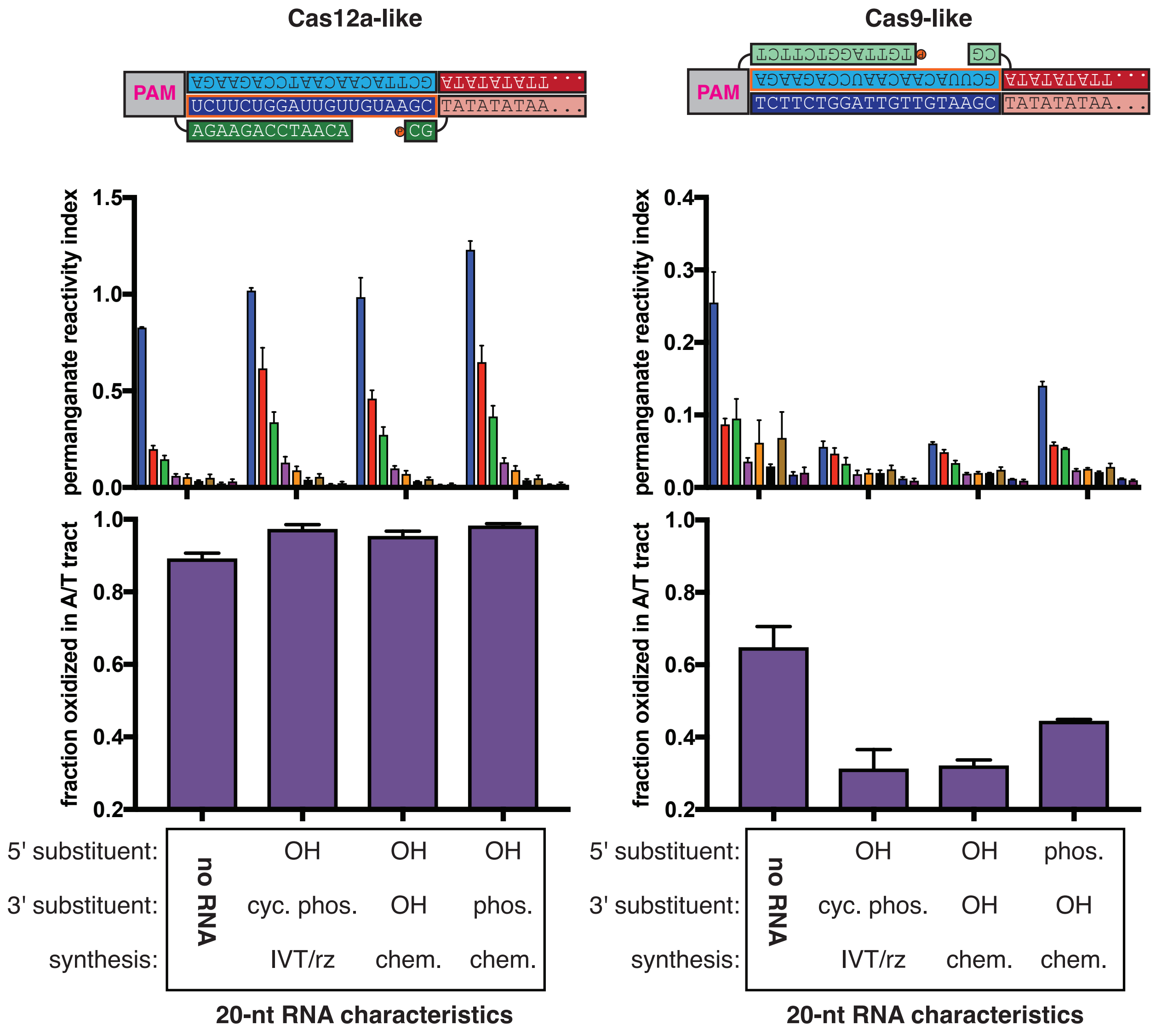


**Supplementary Figure 4D.**

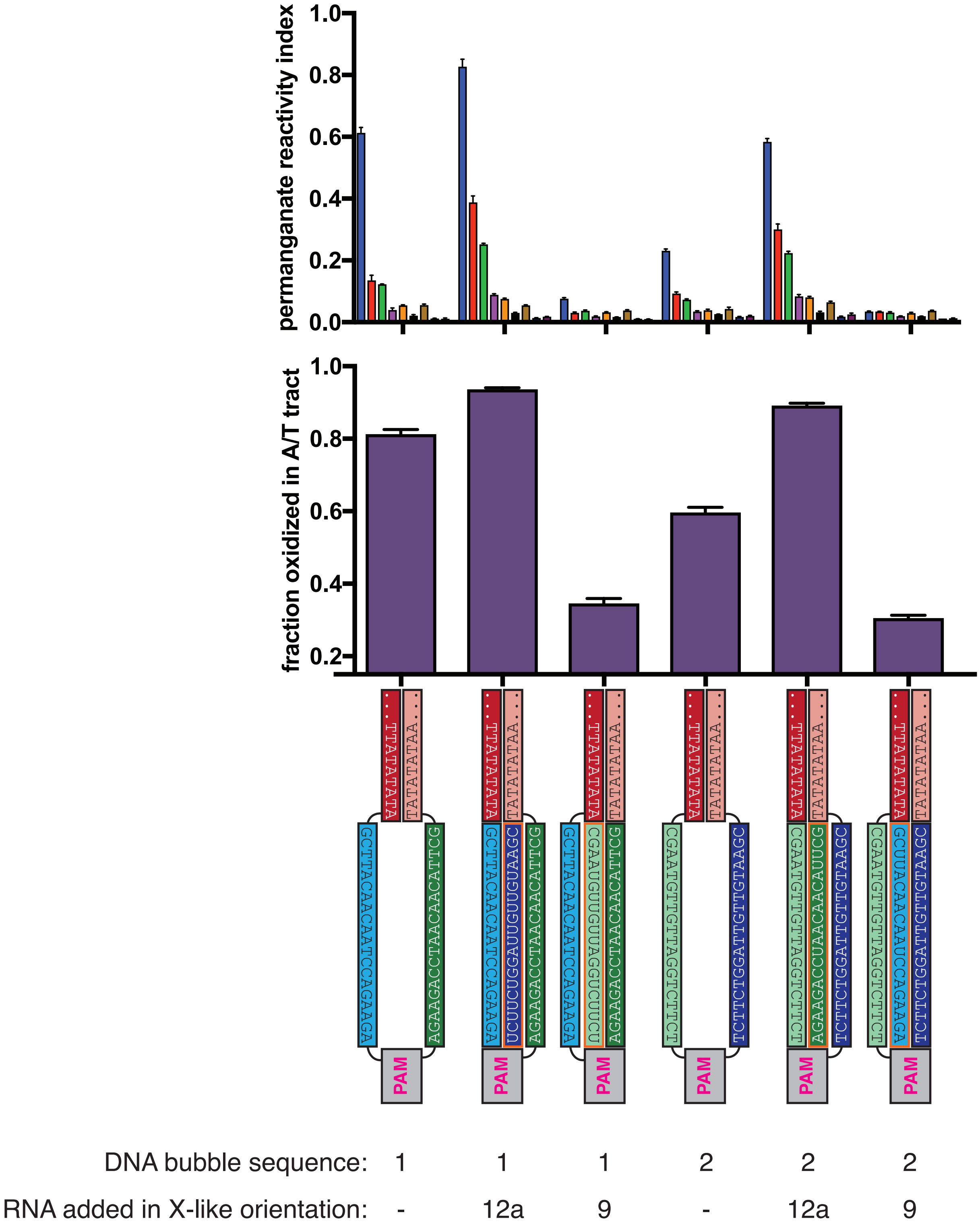


**Supplementary Figure 4E.**

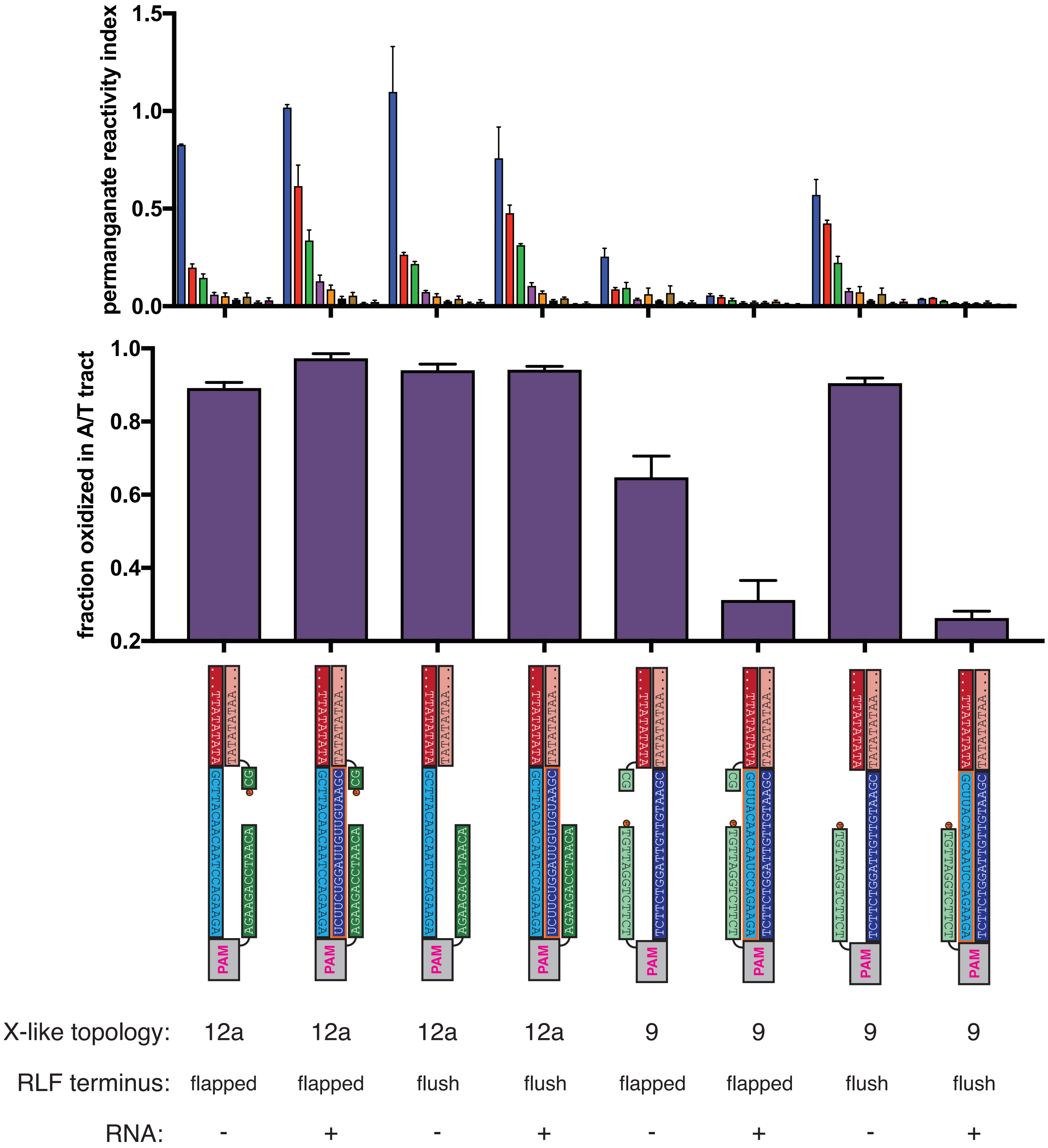


**Supplementary Figure 4F.**

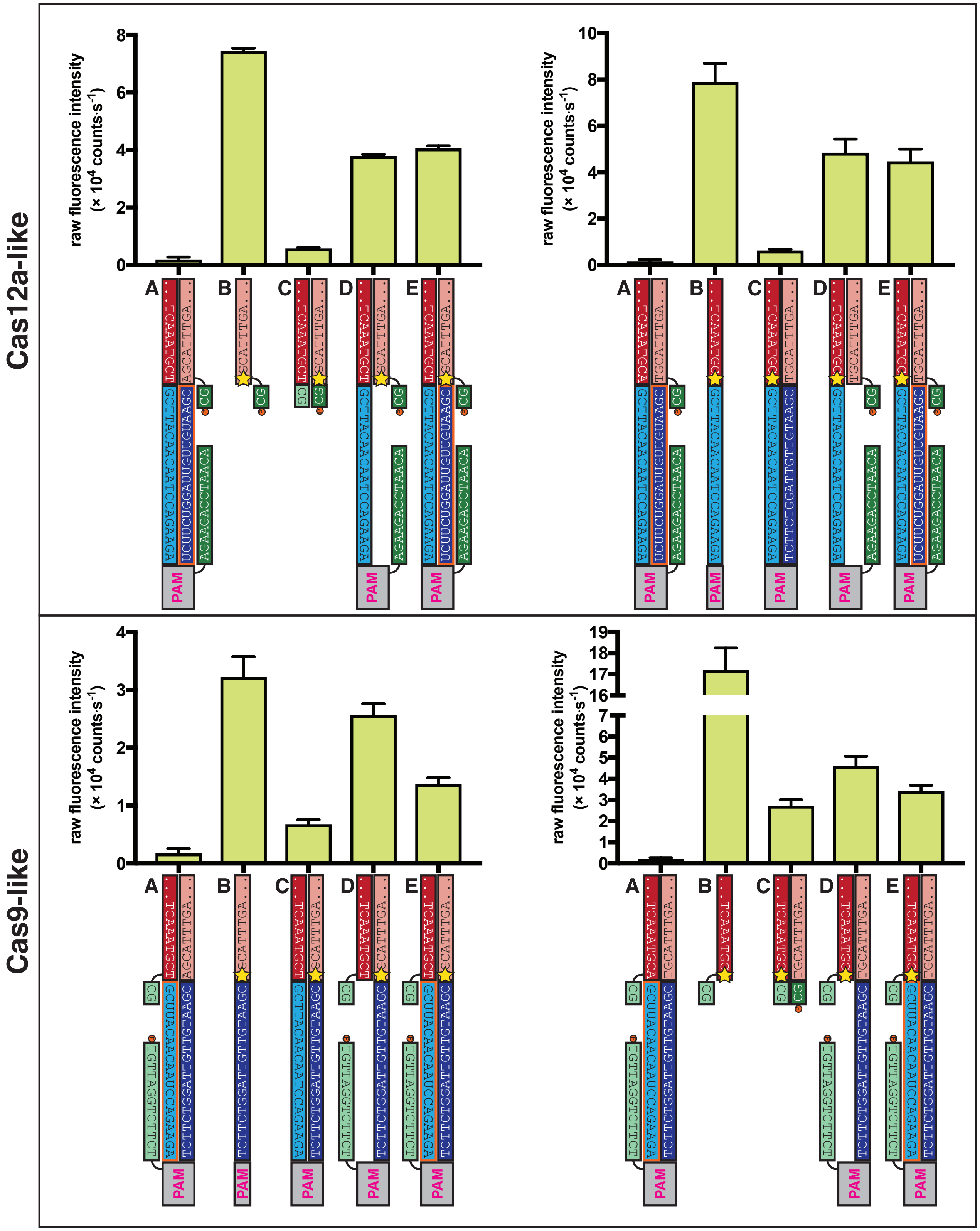


**A:** adenine substituted for 2-aminopurine in **E**
**B:** ssDNA alone
**C:** perfect DNA duplex
**D:** pre-gapped, pre-unwound DNA bubble
**E:** pre-gapped, pre-unwound DNA bubble + 20-nt RNA
★ 2-aminopurine

**Supplementary Figure 4G.**

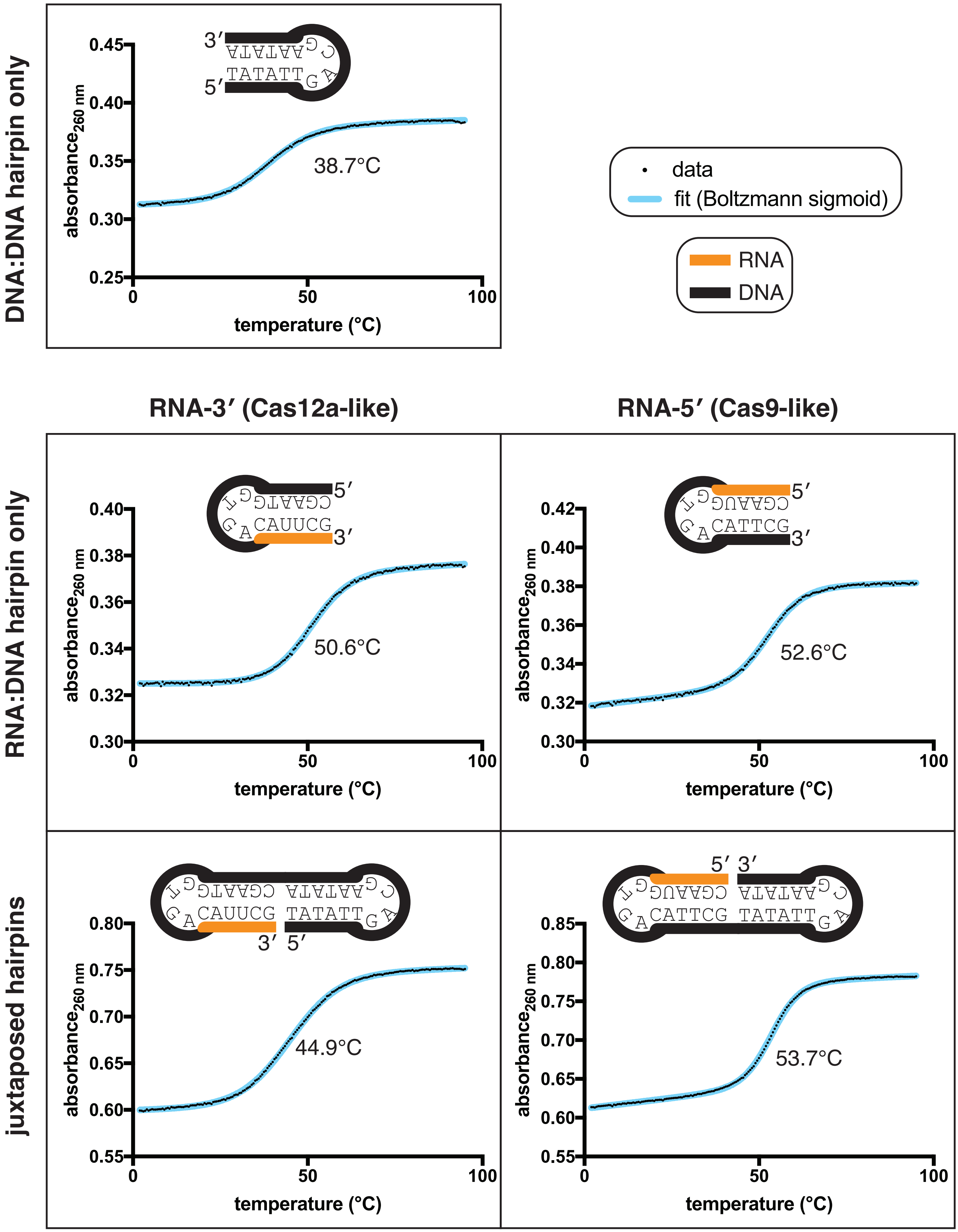


**Supplementary Figure 5A.**

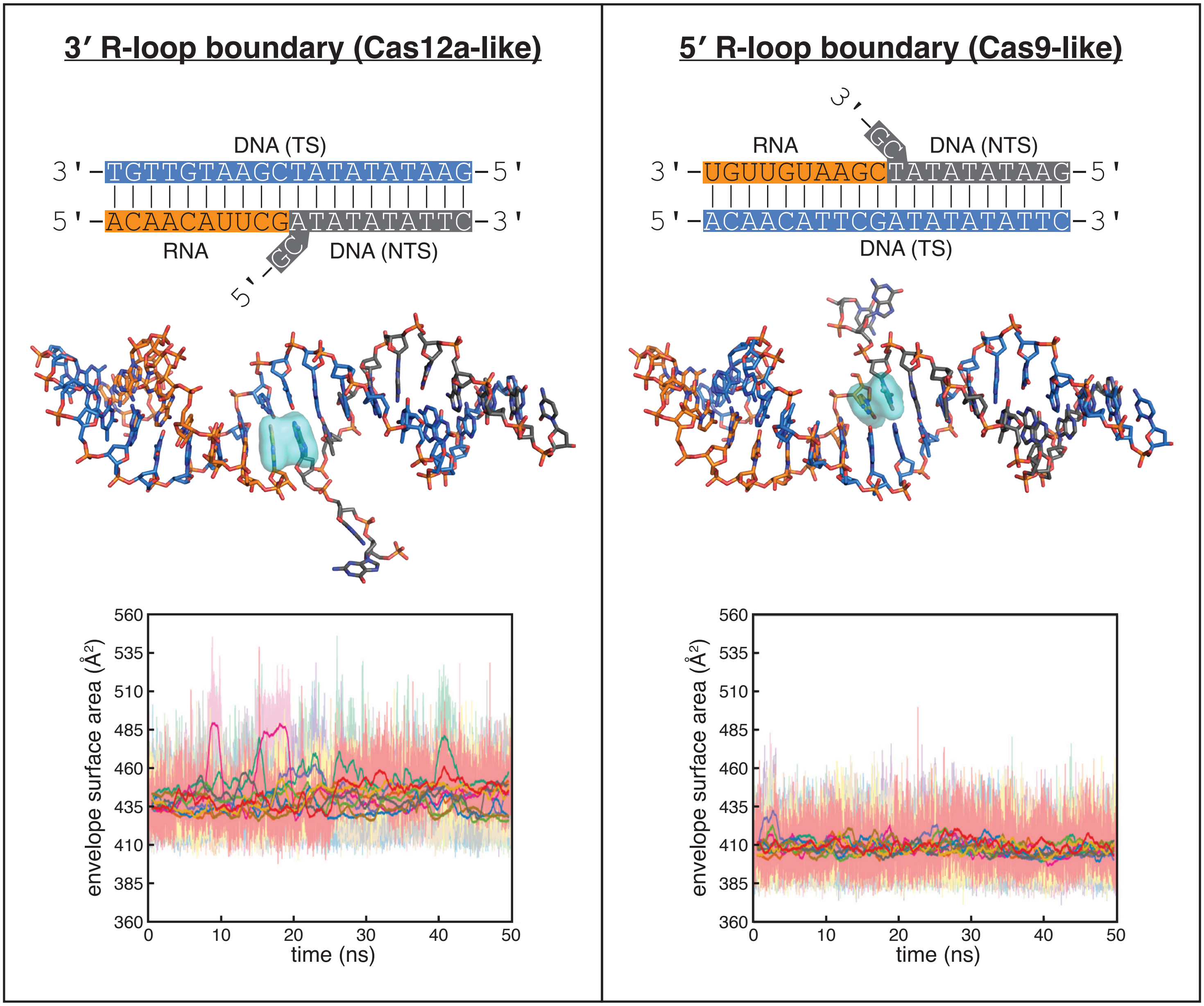


**Supplementary Figure 5B.**

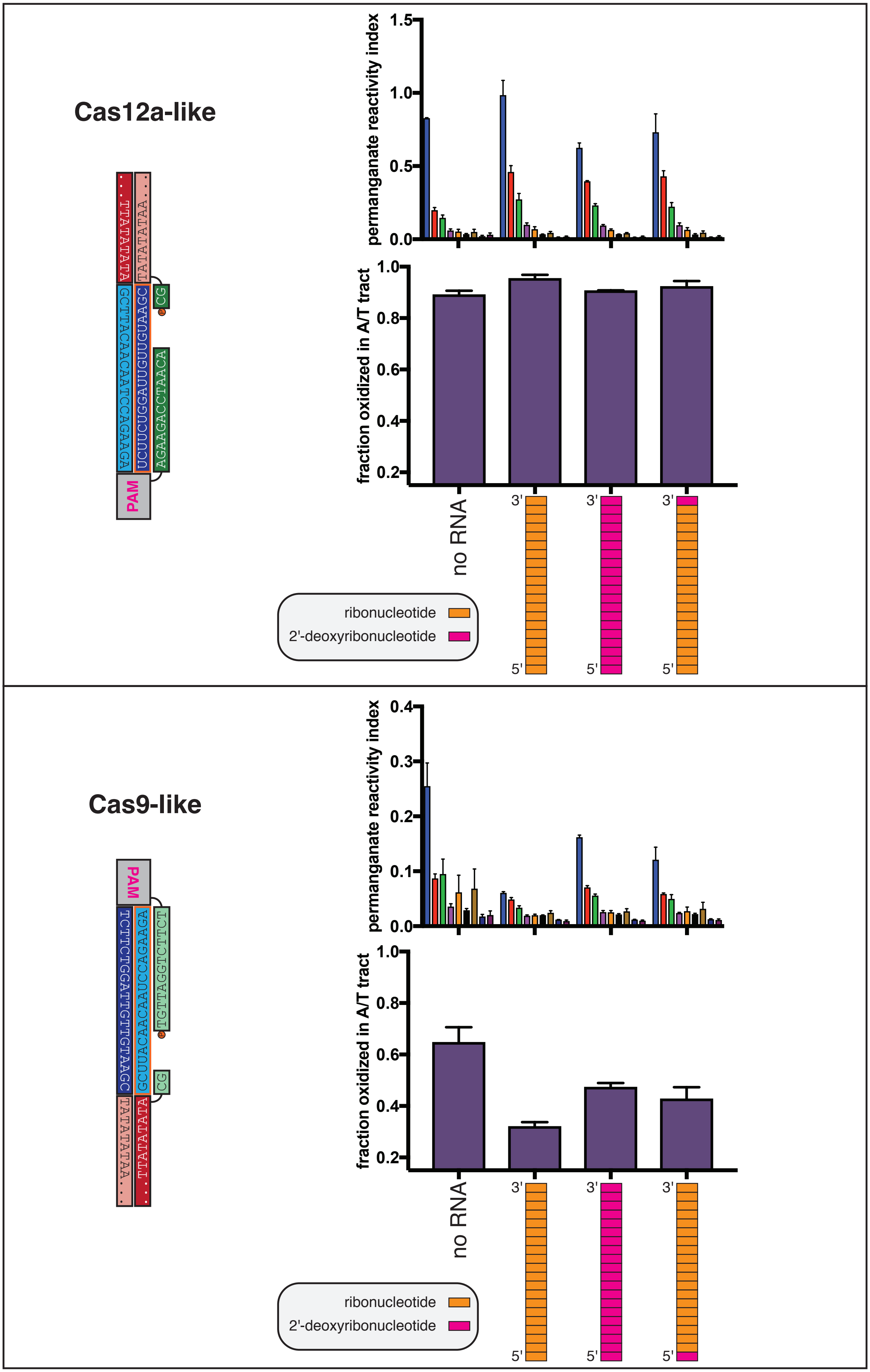


**Supplementary Figure 5C.**

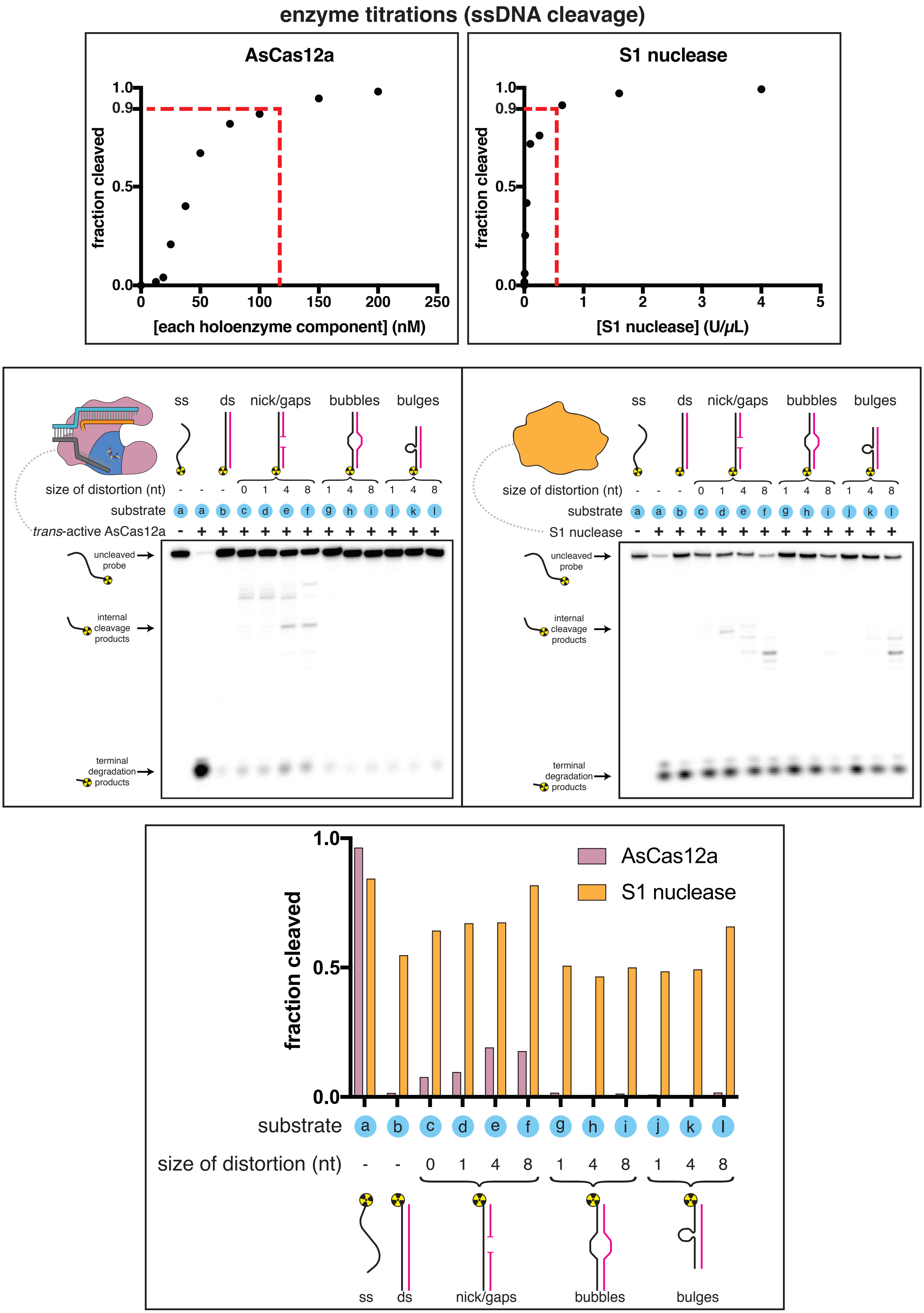


**Supplementary Figure 7A.**

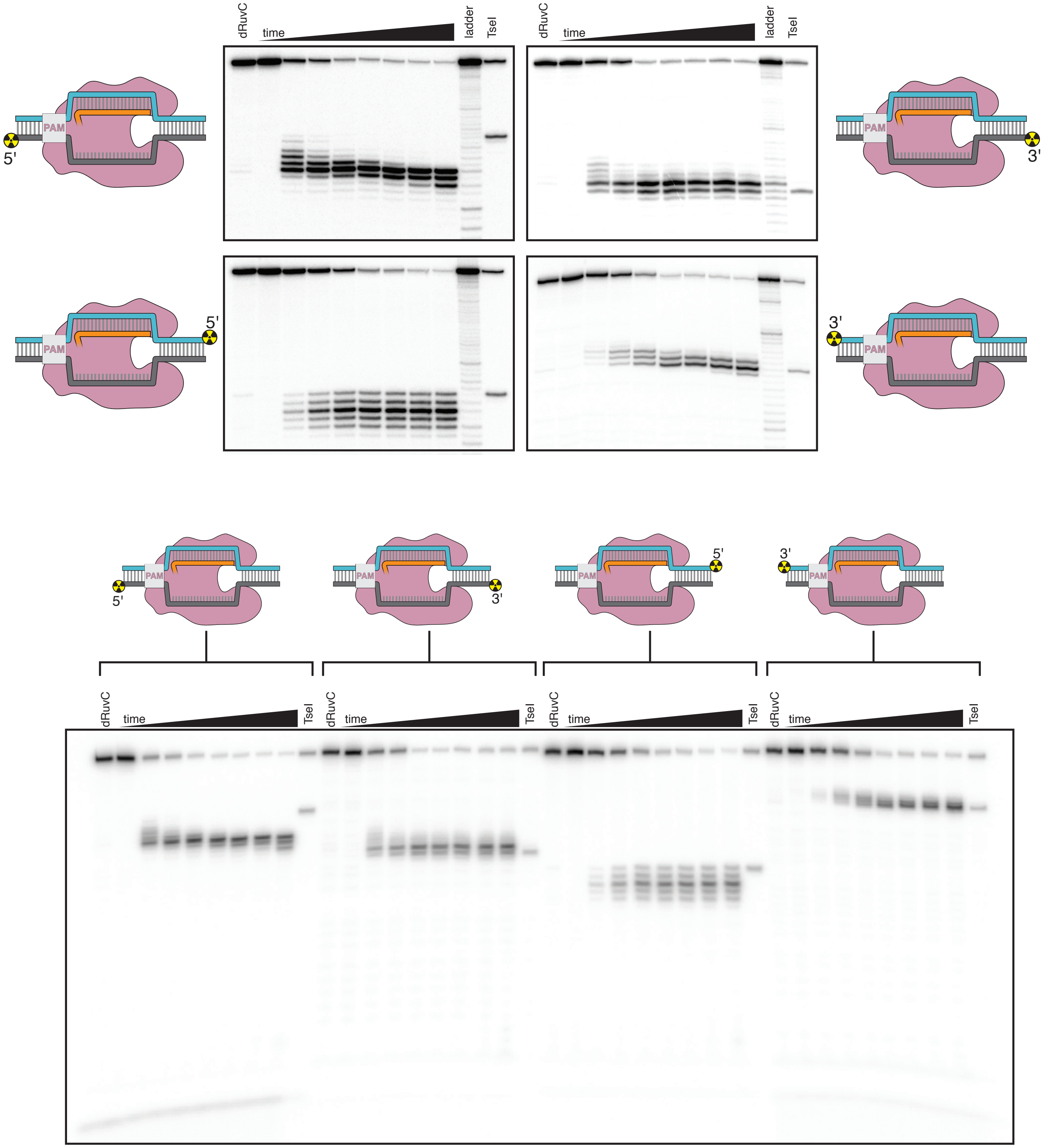


**Supplementary Figure 7B.**

**steady-state kinetics of *trans* ssDNA cleavage, AsCas12a**

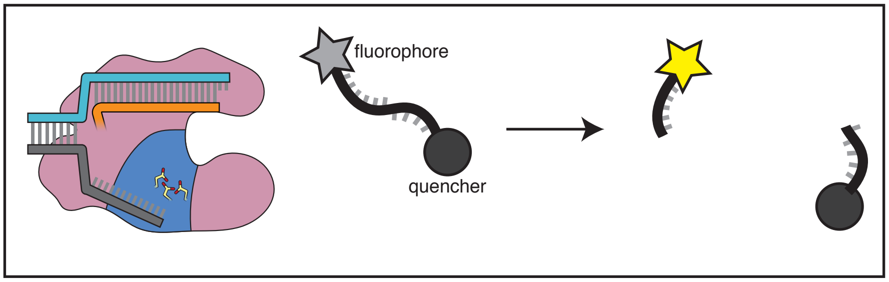


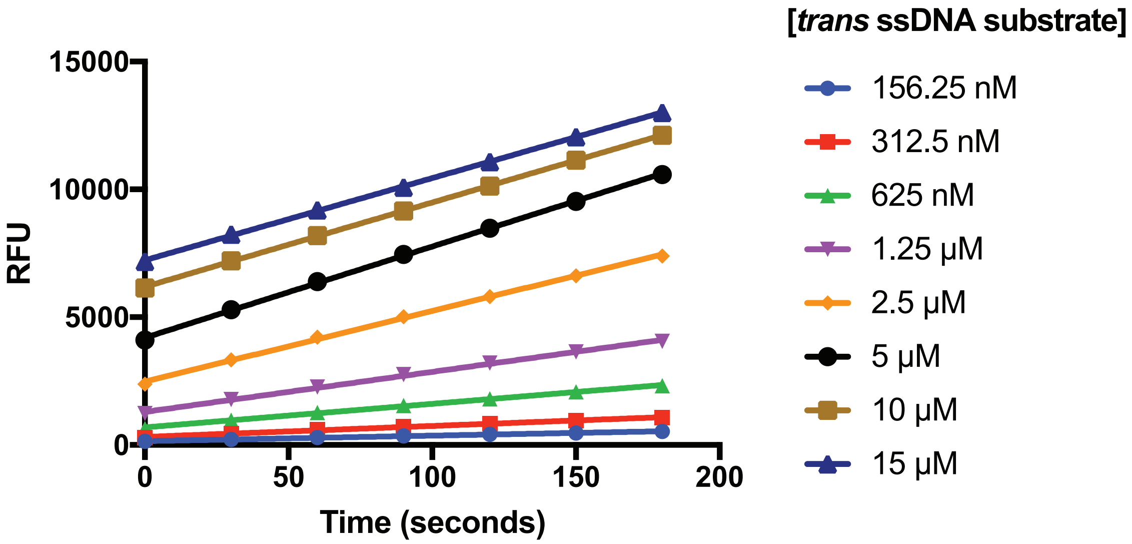


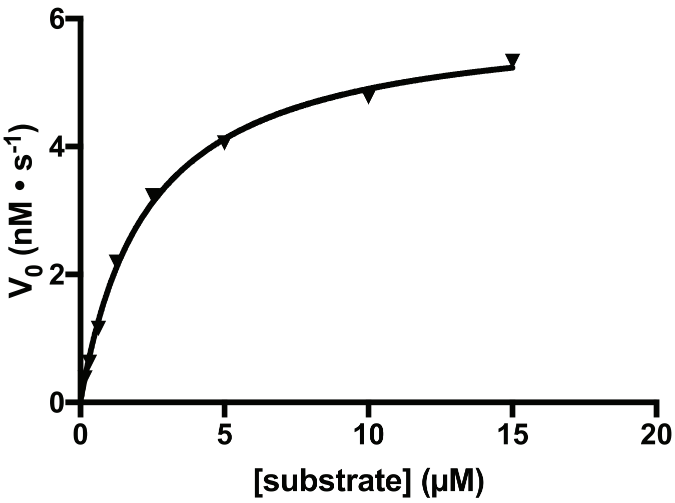


| | |
|---|---|
| $V_{max}$ | 6 nM • $s^{-1}$ ± 1 |
| $K_M$ | 2.7 $\mu$M ± 0.3 |
| $k_{cat}$ | .6 $s^{-1}$ ± 0.1 |
| $k_{cat}/K_M$ | 2.4 × $10^5$ $M^{-1}$ • $s^{-1}$ ± 0.3 |

**Supplementary Figure 7C.**

#### single-turnover *cis* substrate cleavage kinetics, AsCas12a

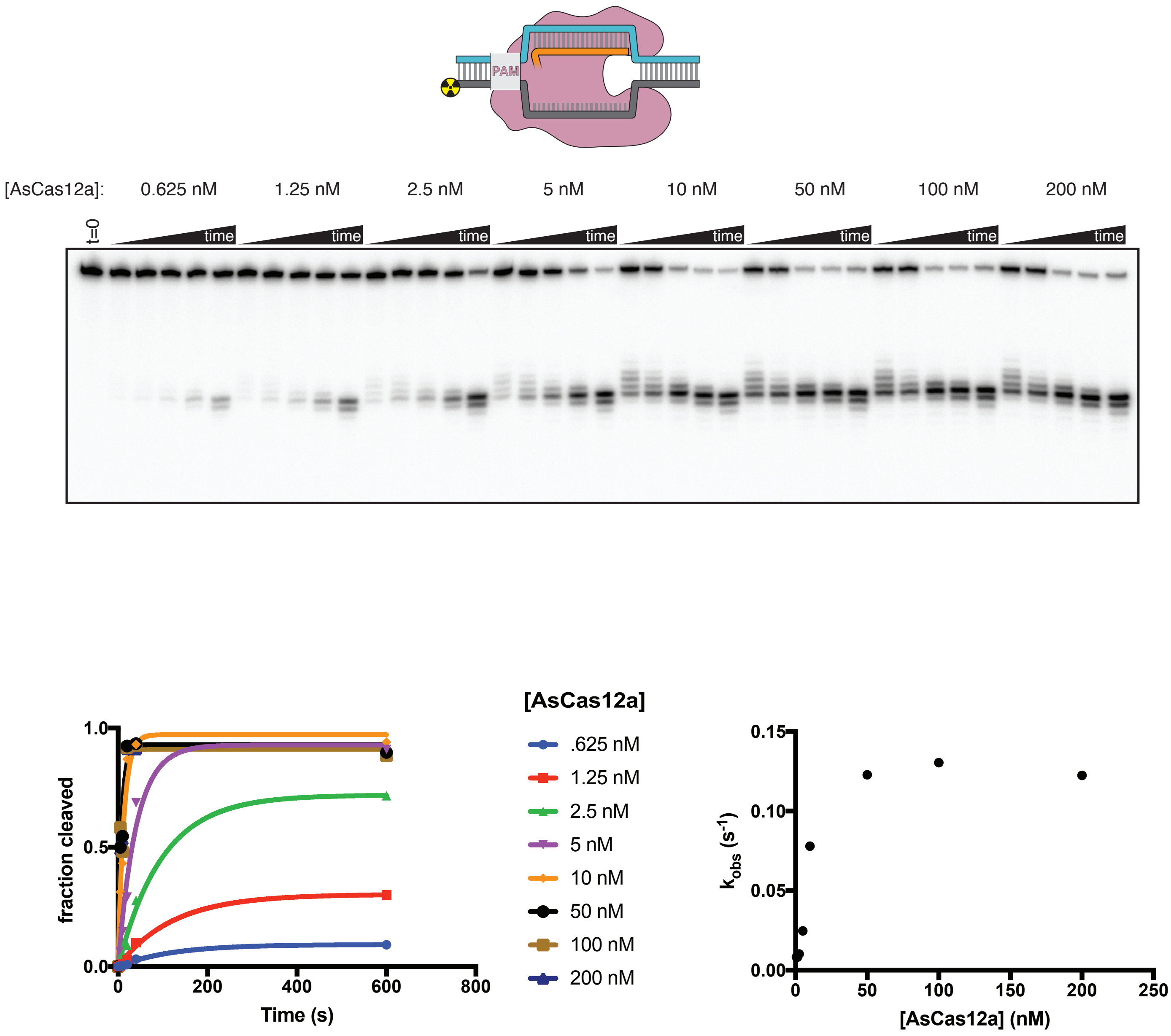


**Supplementary Figure 7D.**

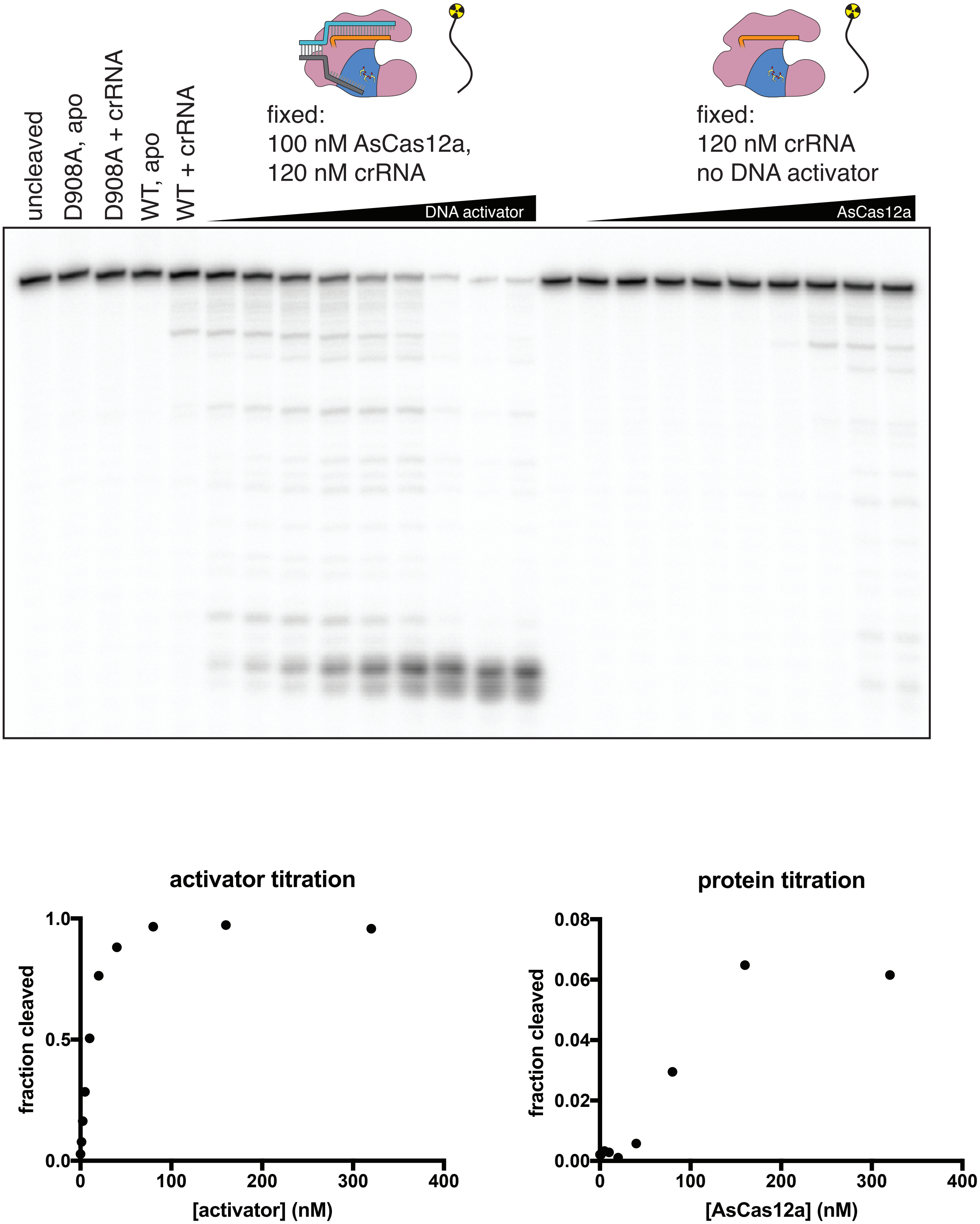


**Supplementary Figure 7E.**

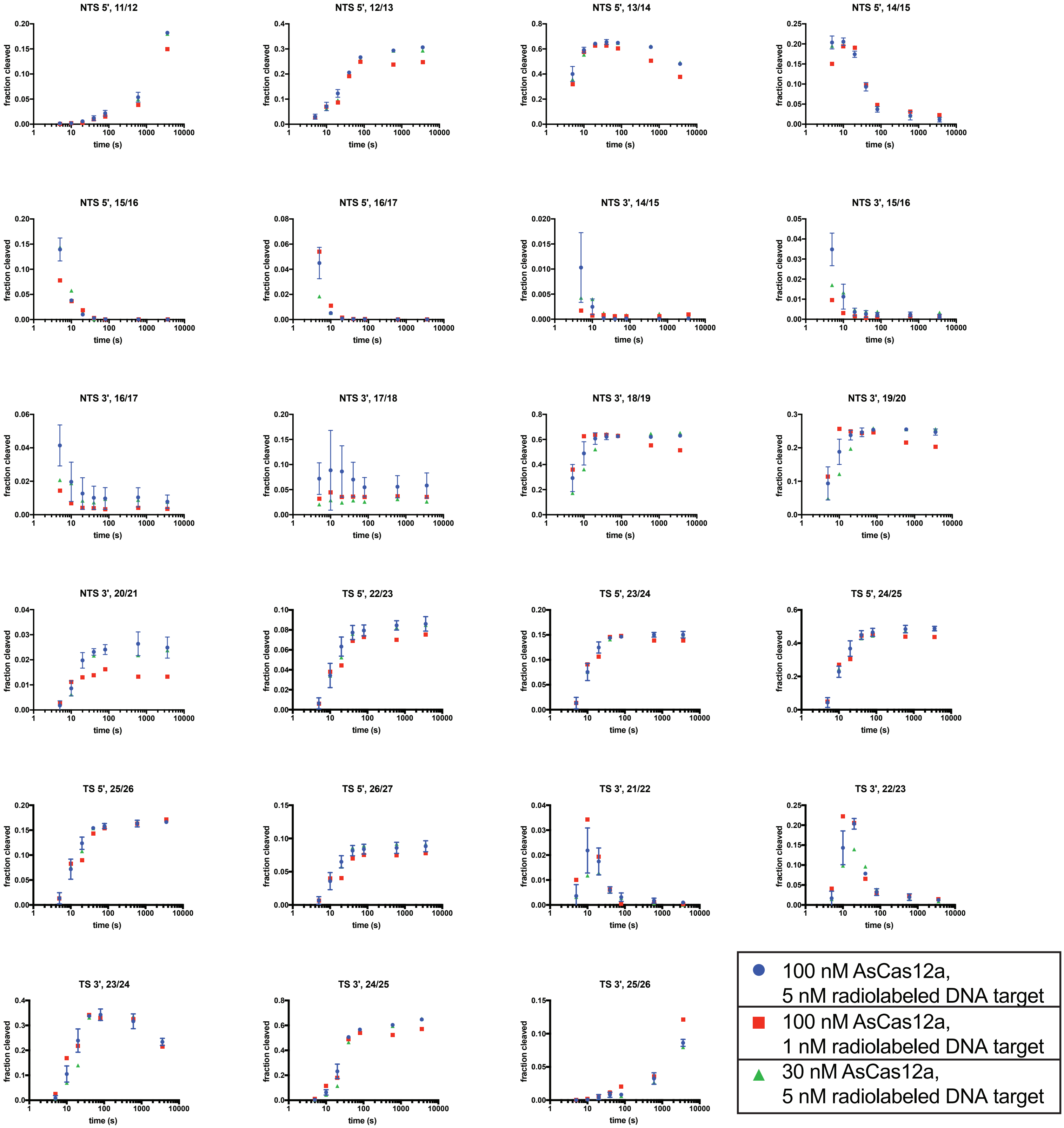


**Supplementary Figure 7F.**

#### cleavage site mapping for FnCas12a

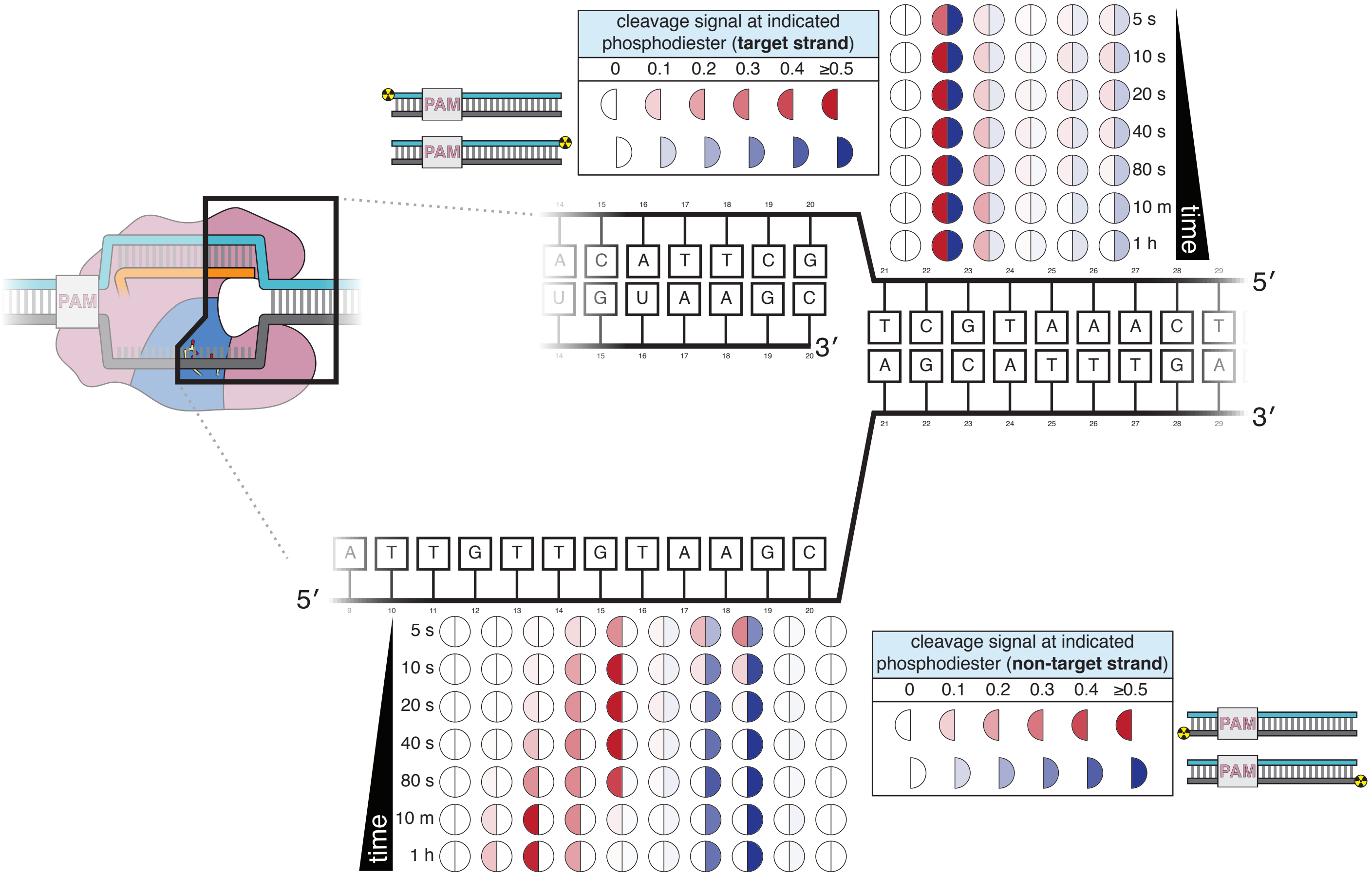


Supplementary Figure 7G.

#### interdependence of NTS cleavage and TS cleavage

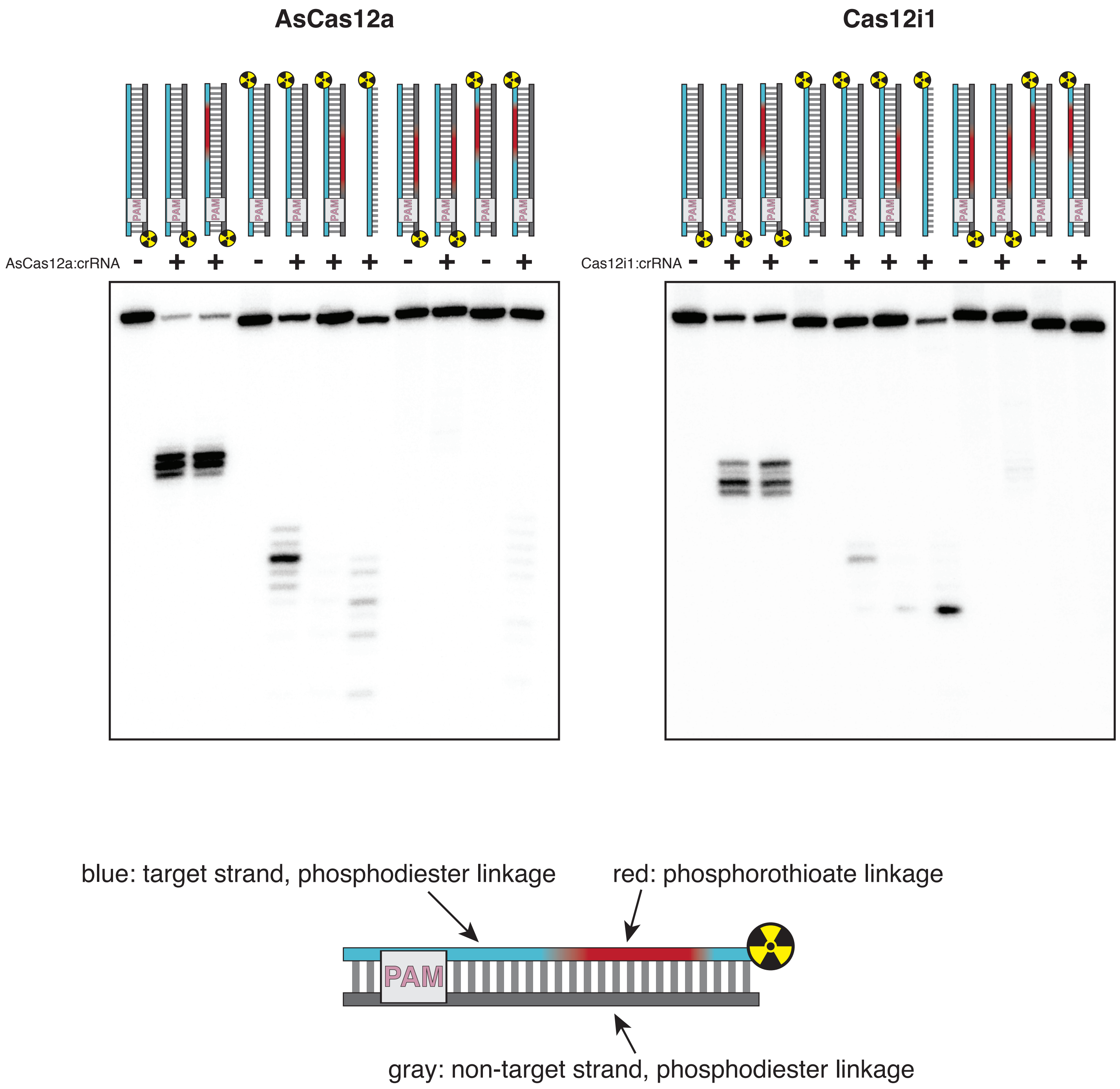


**Supplementary Figure 8A.**

### $MgCl_2$

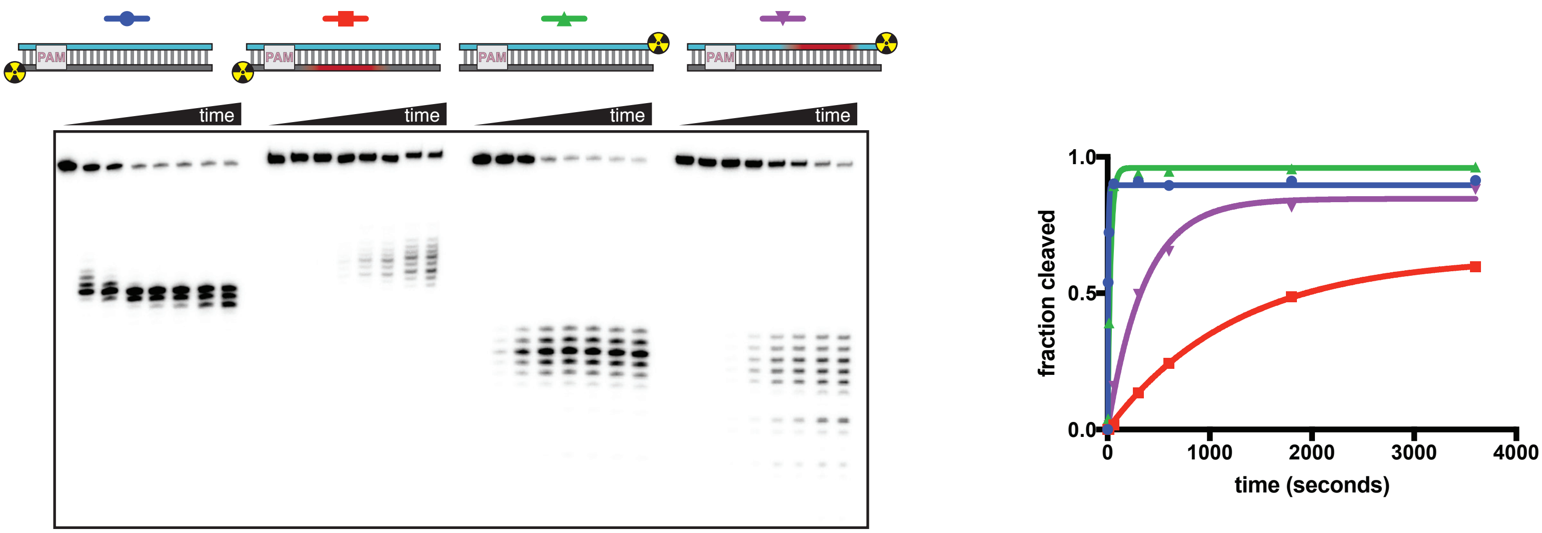


### $CaCl_2$

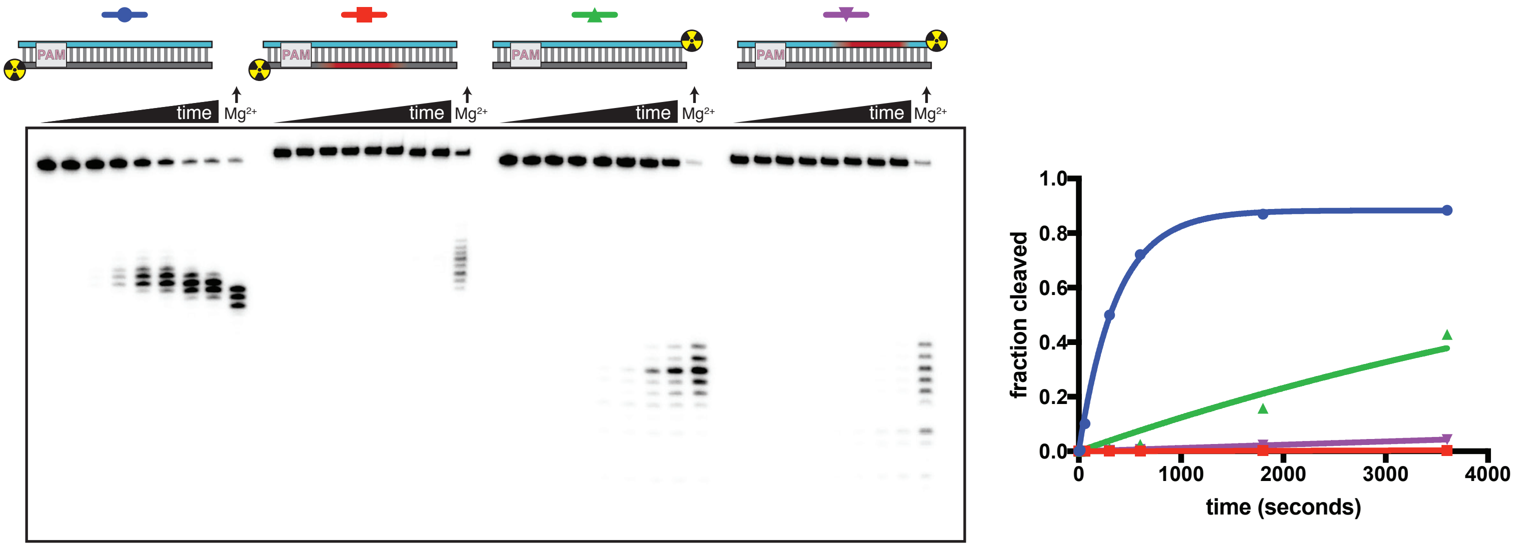


| | $k_{obs}$ ($s^{-1}$), $MgCl_2$ | $k_{obs}$ ($s^{-1}$), $CaCl_2$ | fold change (decrease) |
|---|---|---|---|
| ● | 0.16 (0.11 to 0.22) | 0.0027 (0.0024 to 0.0031) | 57 |
| ■ | $8.0 \times 10^{-4}$ ($7.4$ to $8.7 \times 10^{-4}$) | $1.1 \times 10^{-6}$ ($0.7$ to $1.5 \times 10^{-6}$) | 730 |
| ▲ | 0.034 (0.025 to 0.047) | $1.3 \times 10^{-4}$ ($1.0$ to $1.6 \times 10^{-4}$) | 260 |
| ▼ | 0.0028 (0.0023 to 0.0034) | $1.23 \times 10^{-5}$ ($1.19$ to $1.27 \times 10^{-5}$) | 230 |

**Supplementary Figure 8B.**

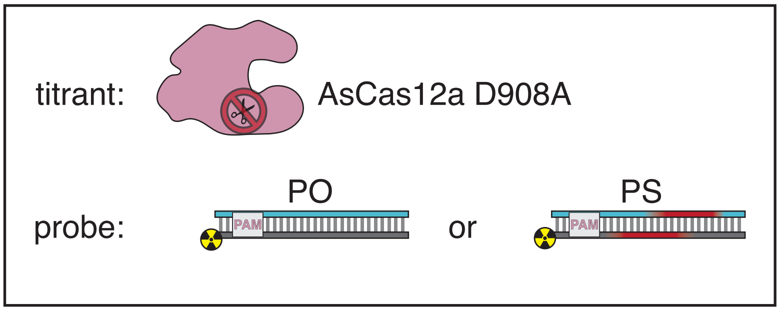


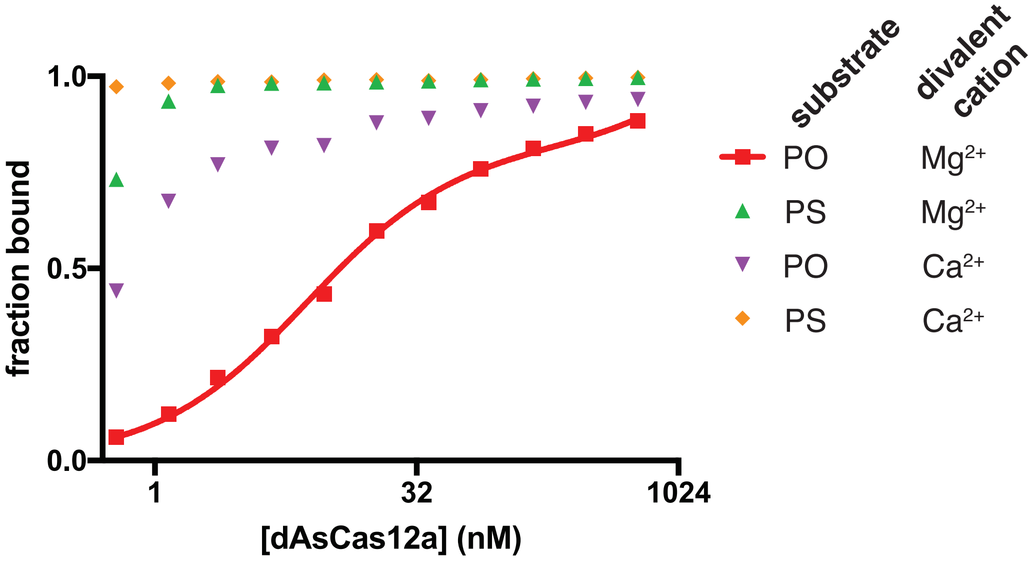


**Supplementary Figure 8C.**

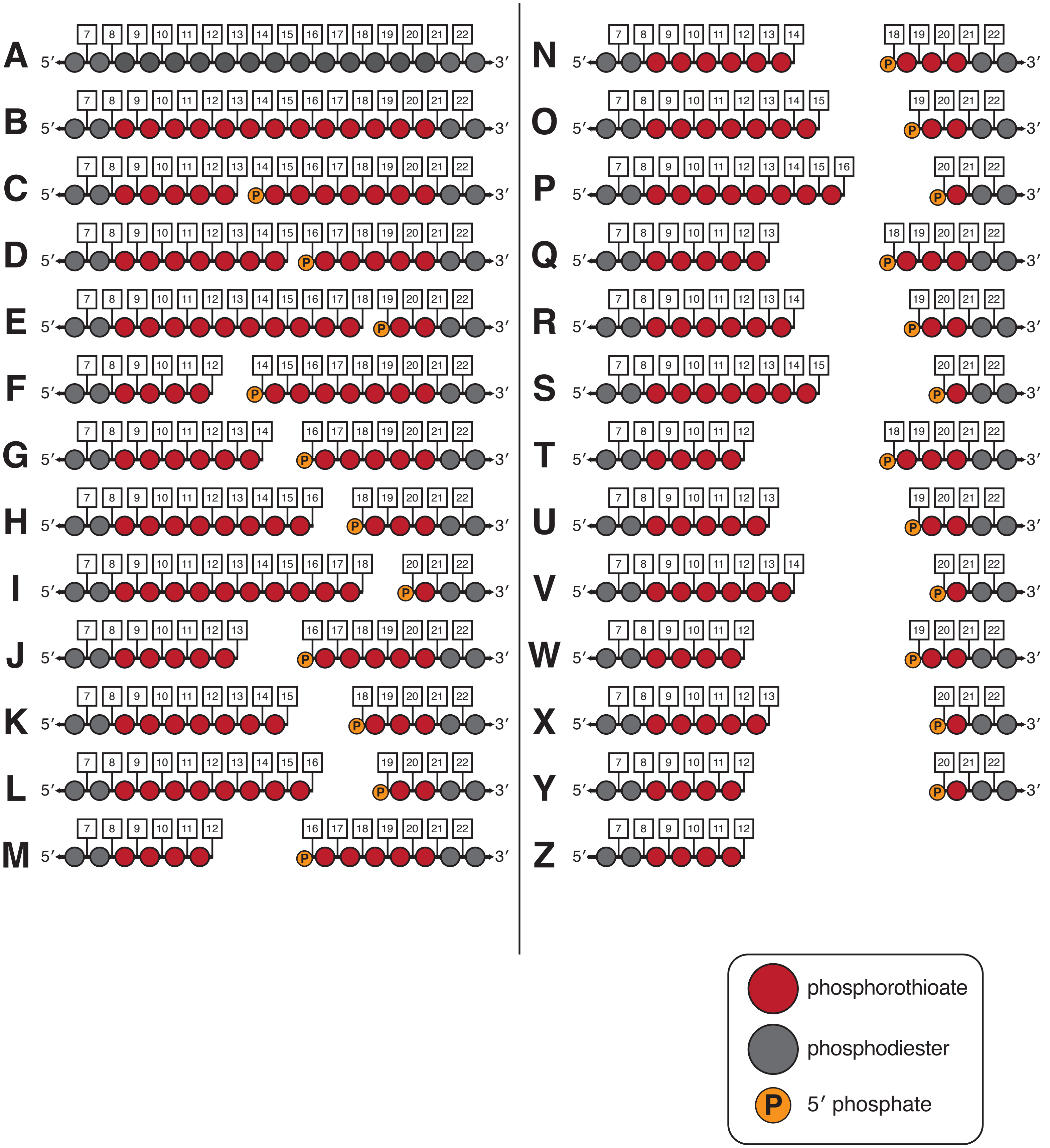


**Supplementary Figure 8D.**

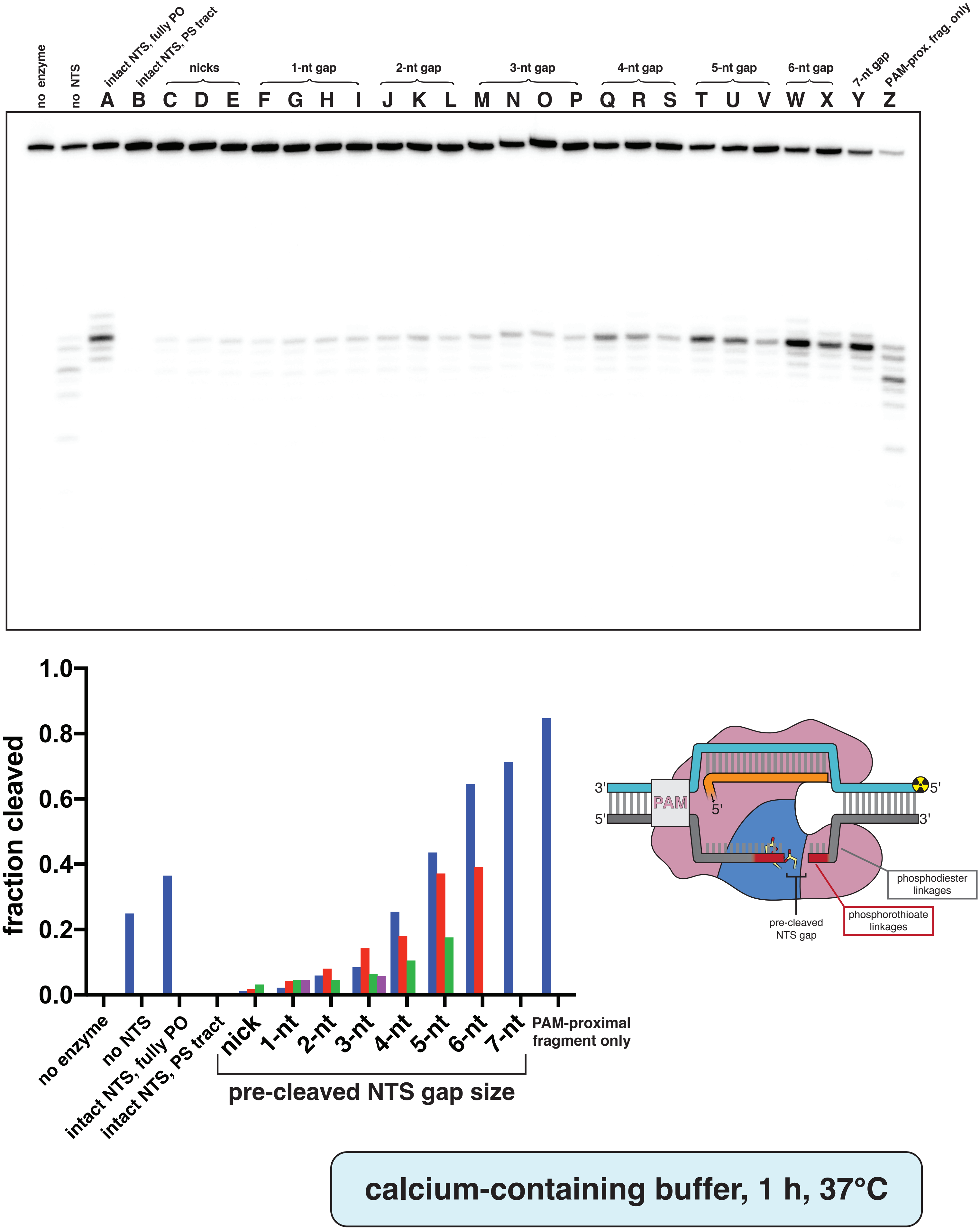


Supplementary Figure 8E.

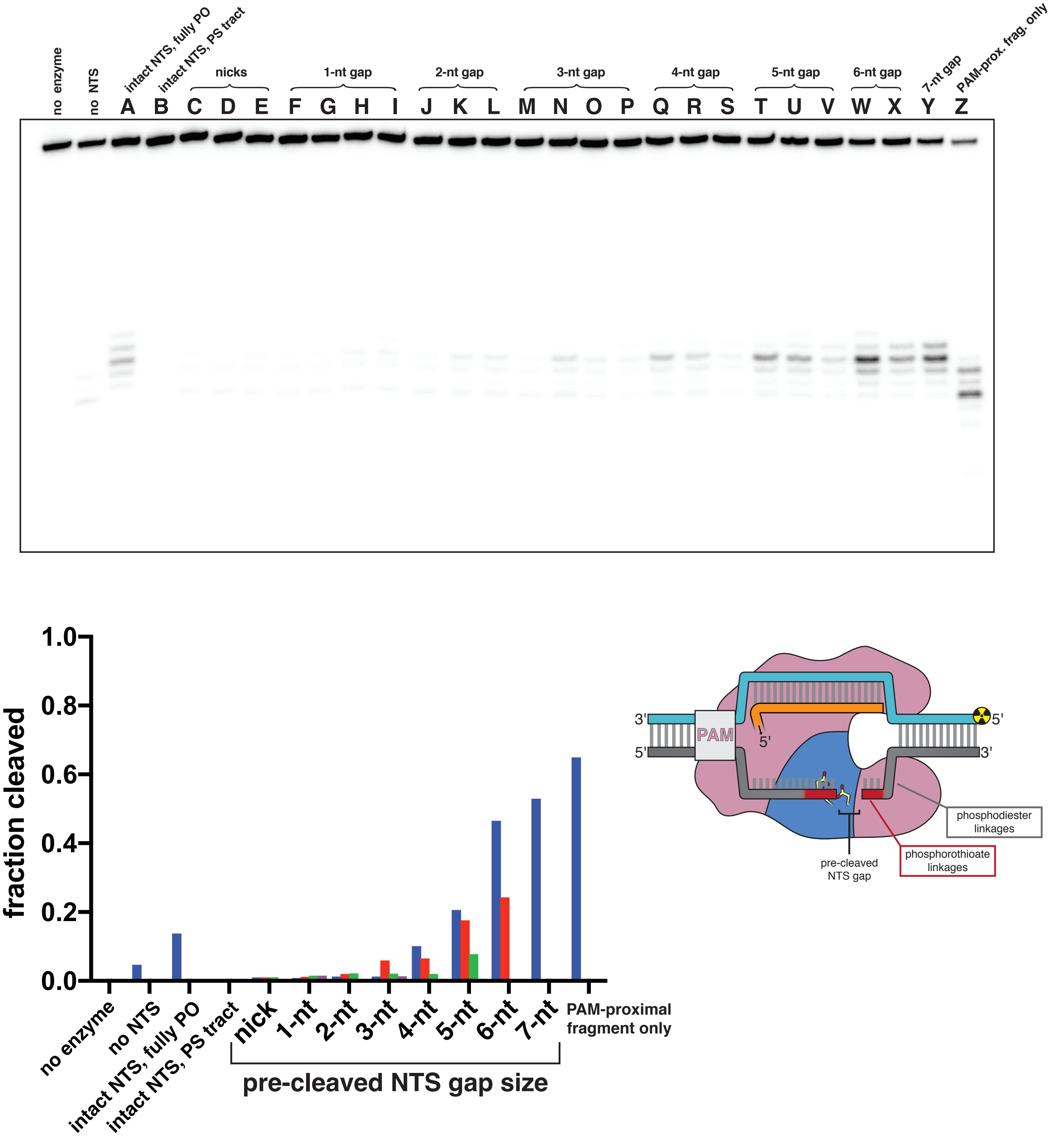


**magnesium-containing buffer, 10 s, 37°C**

Supplementary Figure 8F.

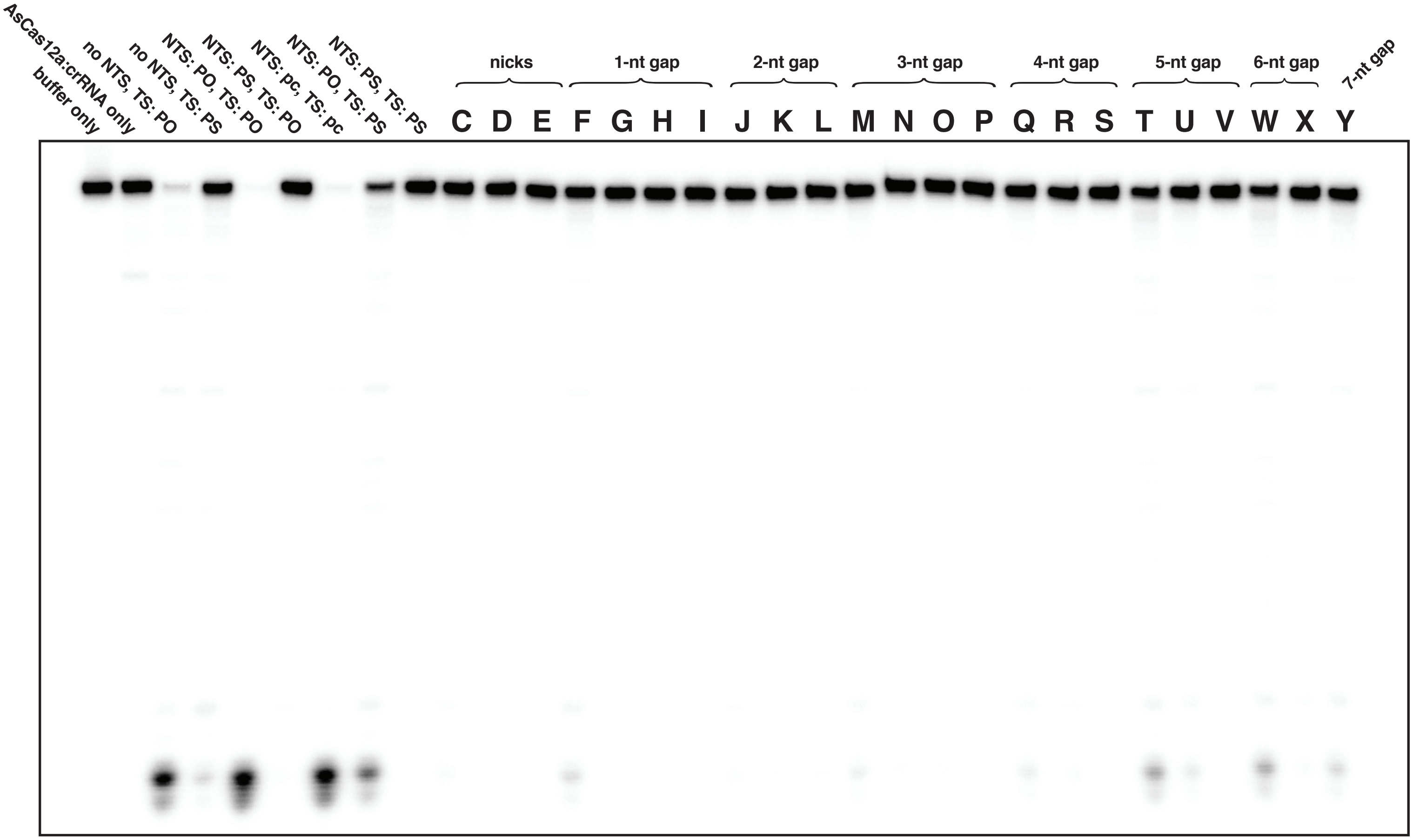


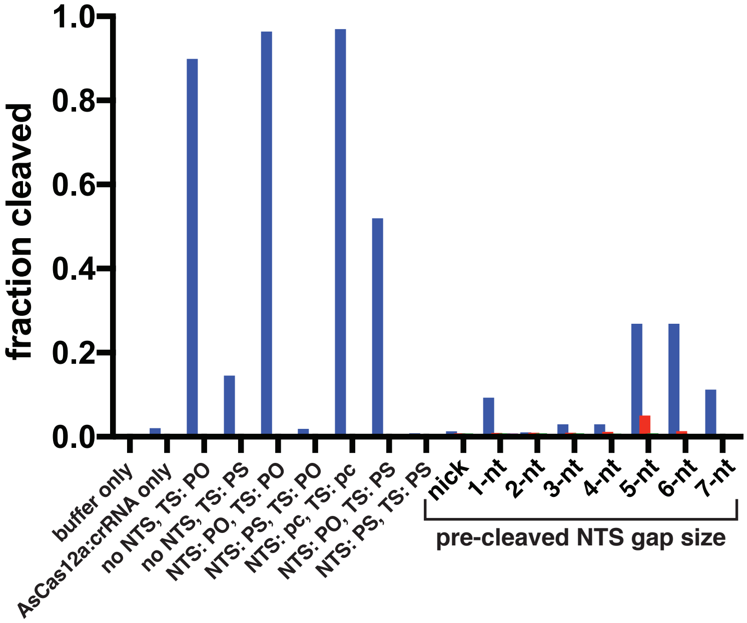


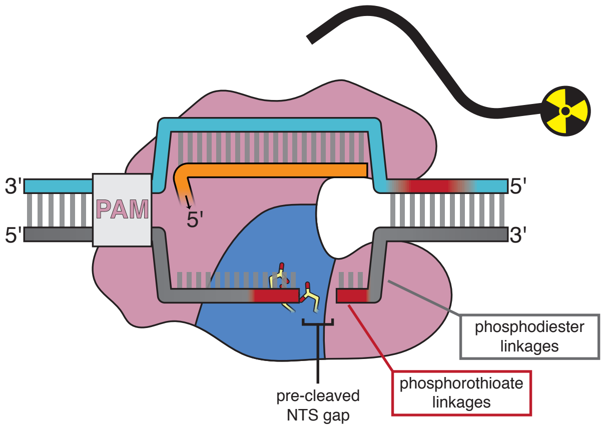


**magnesium-containing buffer, 30 m, 37°C**

Supplementary Figure 8G.

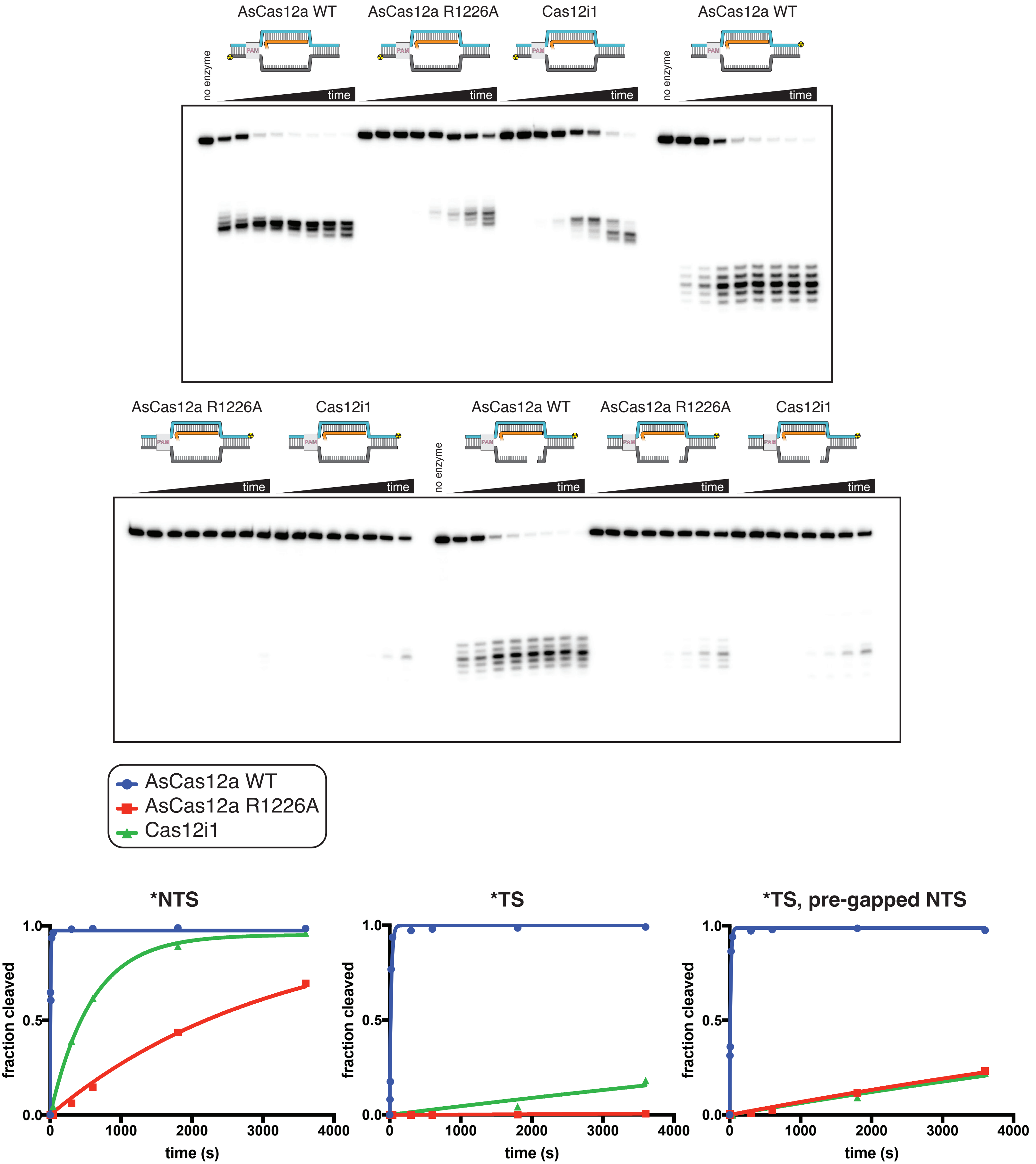


**Supplementary Figure 8H.**

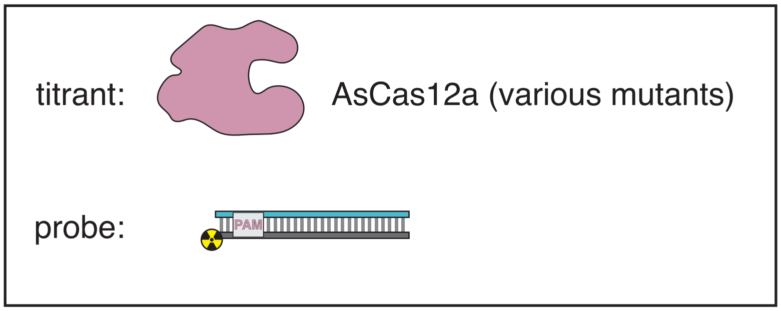


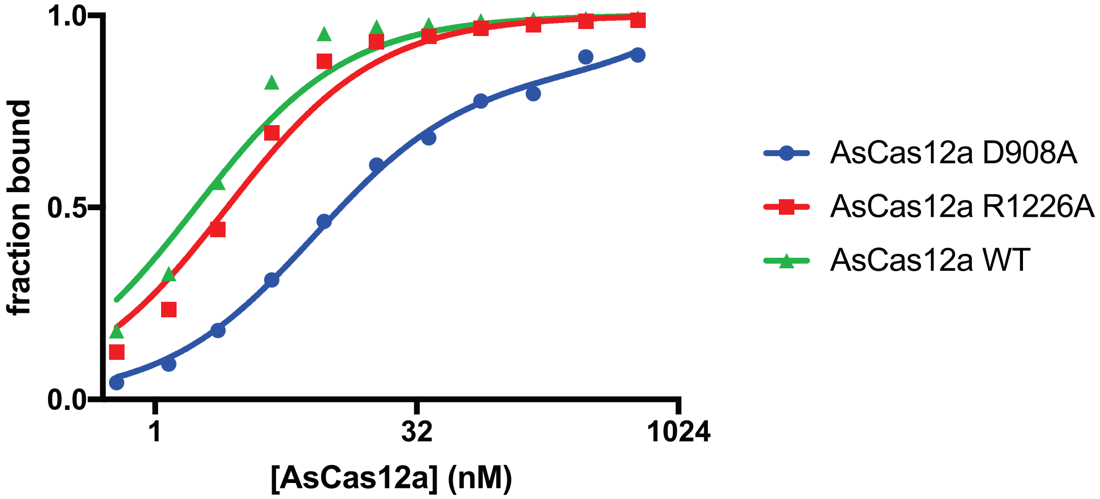


**Supplementary Figure 8I.**

cleavage site mapping for AsCas12a R1226A

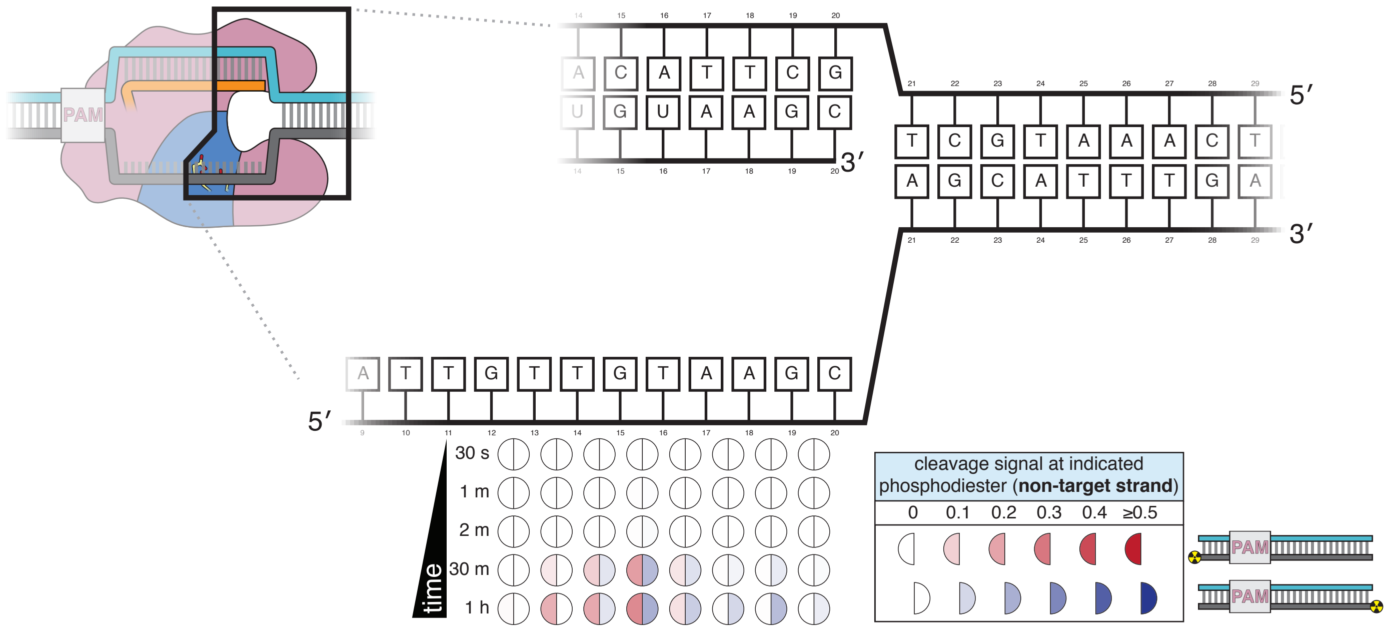


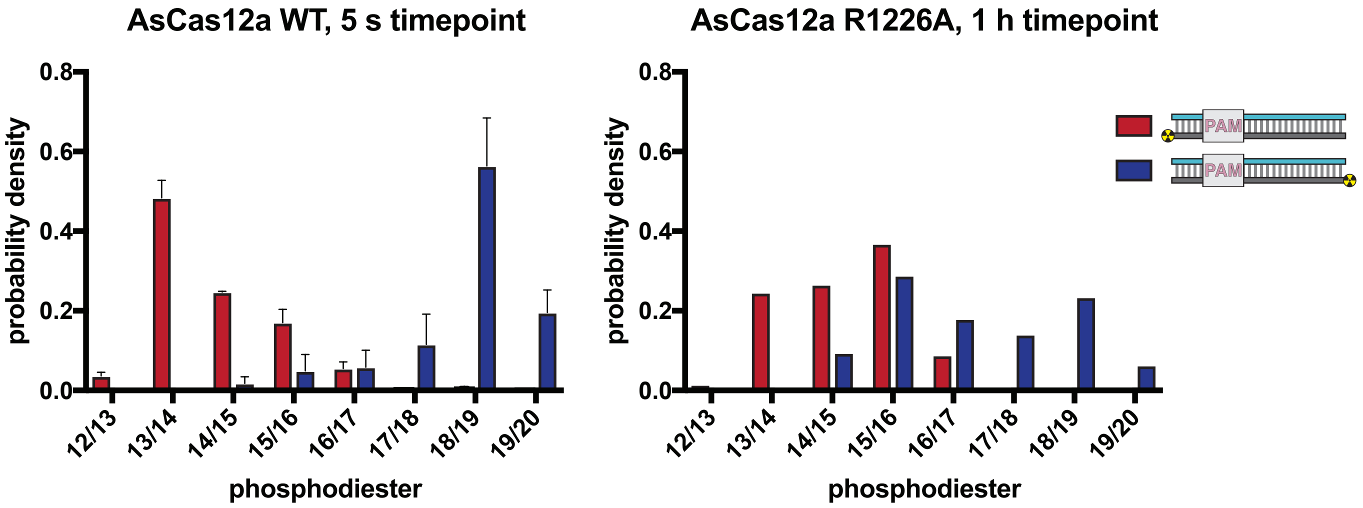


Supplementary Figure 8J.

#### cleavage site mapping for Cas12i1

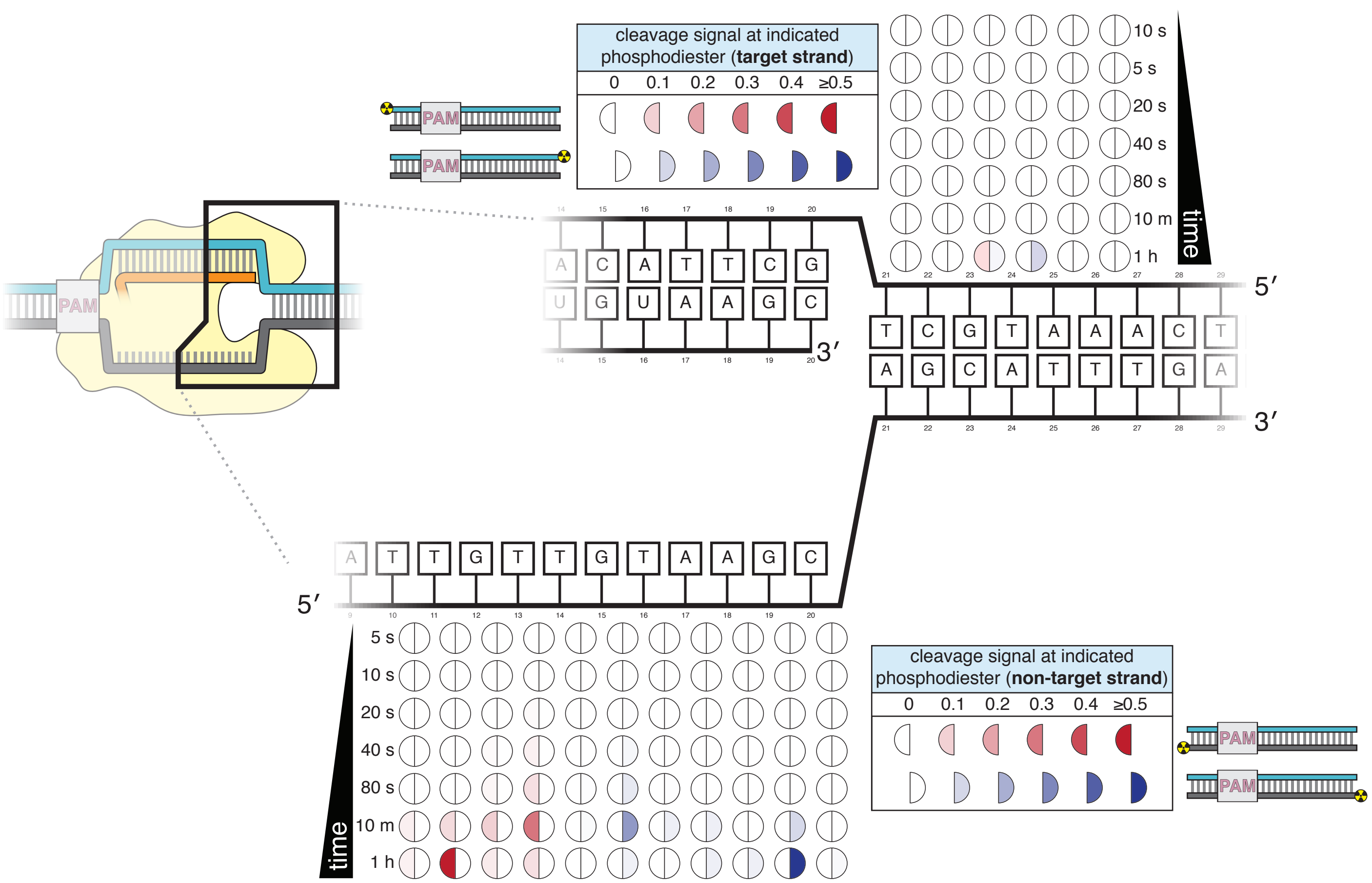


Supplementary Figure 8K.

#### SUPPLEMENTARY FIGURE LEGENDS

**Supplementary Figure 2A. Method used to 3′-end radiolabel DNA oligonucleotides.** See Methods section for details.

**Supplementary Figure 2B. A gap in the non-target strand increases the affinity of dCas12a for its DNA target.** *Top panel*: The affinity of dAsCas12a/crRNA for a cognate DNA target was assessed by an electrophoretic mobility shift assay (EMSA) and a filter-binding (FB) assay. dAsCas12a was titrated in a solution with fixed [crRNA] (750 nM) and [DNA probe] (100 pM), followed by separation of protein-bound DNA from free DNA. The EMSA indicated that the oligonucleotide annealing protocol yields 100% duplex DNA probe and that the binding conditions yield one major protein-bound species. “Fraction bound” is defined as (background-subtracted volume of upper band)/(total background-subtracted lane volume) for the EMSA and (background-subtracted volume of Protran spot)/(total background-subtracted volume of Protran spot + Hybond N+ spot) for the filter-binding assay. The value of “fraction bound” was 0 at [dAsCas12a]=0 for both assays (not shown in plot due to the logarithmic x-axis). When appropriate, data were fit to the sum of a hyperbola and a line ($y=B_{max}*x/(K_D+x)+NS*x$), where NS describes a non-specific binding mode. It is common to see $B_{max}$ values below 1 in EMSAs and filter binding assays, in which the process of physical separation can disrupt bound species. $K_D$ for the EMSA was 8.2 nM (n=1). $K_D$ for the filter-binding assay was 8.1 nM ± 0.8 (SD) (n=3). *Bottom panel*: Using the filter binding assay, we assessed the affinity of dAsCas12a/crRNA for various cognate DNA targets. Protospacer 2 (used in **Fig. 2B**) is the version of protospacer 1 (used in **Fig. 2A**) modified for permanganate probing of the R-loop flank. Differences between protospacer 1 and protospacer 2 are highlighted in red (A/T base pairs substituted into the RLF, G/C base pairs substituted elsewhere to maintain stable association between the two DNA strands). “Intact” protospacers are as shown in the sequence schematic. “Pre-gapped” protospacers are missing nt 14-18 of the NTS (as measured from the PAM, see **Appendix B**). The value of “fraction bound” was 0 at [dAsCas12a]=0 for all substrates (not shown due to the logarithmic x-axis). Data were analyzed as described for the top panel. $K_D$ for protospacer 1 (intact) was 5 nM ± 1 (SD) (n=3). $K_D$ for protospacer 2 (intact) was 54 nM ± 12 (SD) (n=3). $K_D$ for protospacer 2 (pre-gapped) was 2.8 nM ± 0.5 (SD) (n=3). Data from the protospacer 1 (pre-gapped) experiment indicated that the $K_D$ was near or below [DNA probe], preventing accurate $K_D$ determination by hyperbolic fitting. The reason for the low observed affinity of dAsCas12a for protospacer 2 (intact) is unknown.

**Supplementary Figure 2C. Translating raw phosphorimages into quantitative permanganate reactivity metrics.** *Left panel*: Kinetics of permanganate reaction with an unpaired thymine. The depicted substrate was subject to the standard permanganate reaction protocol with quenching at 0, 5, 10, 30, 60, and 120 seconds. Black arrows indicate chemical cleavage fragments that resulted from oxidation of the annotated thymine. "Fraction oxidized at thymine X," plotted in the graph at the bottom, was determined as described in **Supp. Note 1** and is equivalent to the variable $p_i$. The phosphorimage and graph shown are from a single representative replicate (n=3). Data were fit to an exponential decay (y=($y_0$-plateau)*exp(-k*x)+plateau), with $y_0$ constrained to 0 and the plateau value constrained to 1. The value of k was determined to be 0.998 $min^{-1}$ ± 0.027 (SD) (n=3), which, when corrected to the reference thymine, yielded the value of $k_{ss,corr}$=0.79 $min^{-1}$ that was used for normalization of the permanganate reactivity index in all other permanganate experiments. *Right panel*: Raw phosphorimage of quantified data presented in **Fig. 2B**. Black arrows indicate chemical cleavage fragments that resulted from oxidation of the annotated thymine. The method to determine the "permanganate reactivity index" and "fraction oxidized" metrics from a raw phosphorimage is described in **Supp. Note 1**.

**Supplementary Figure 3A. dCas12a ribonucleoprotein binds tightly to pre-gapped/pre-unwound targets despite PAM-distal mismatches.** The affinity of dAsCas12a/crRNA for various cognate DNA targets was assessed by a filter-binding assay. “Pre-gapped” indicates the presence of a 5-nt gap in the non-target strand (see **Appendix B**). “Pre-unwound” indicates the presence of a stretch of NTS:TS mismatches in the DNA substrate. In **Fig. 3A**, protospacer 3 is annotated as “DNA substrate 1;” protospacer 4 is annotated as “DNA substrate 2;” and crRNA 3 is the depicted crRNA. For each combination of crRNA/DNA target, crRNA was titrated in a solution with fixed [dAsCas12a] (400 nM), [DNA probe] (100 pM), and [non-specific DNA competitor] (500 nM). The identities of the titrant/fixed component were inverted in this experiment (as compared to all other binding experiments) because crRNA can form a stable complex with pre-unwound DNA targets in the absence of protein. Keeping [dAsCas12a] at 400 nM favored the formation of (dAsCas12a/crRNA):DNA complexes over crRNA:DNA complexes (which would be indistinguishable from free DNA in the filter binding assay). In the presence of high [apo protein], 500 nM non-specific DNA competitor (a duplex with a short ssDNA overhang) was also included to disfavor non-specific interactions between radiolabeled DNA and apo protein. The value of “fraction bound” was 0 at [crRNA]=0 for all substrates (not shown due to the logarithmic x-axis). For all pre-unwound DNA targets, the fraction bound was essentially concentration-independent across all nonzero concentrations tested, suggesting that the lowest concentration tested had already saturated the specific binding interaction being probed. The high stability is in line with thermodynamic expectations for an interaction involving hybridization of two complementary 18-nt or 20-nt oligonucleotides ($T_m$ > 40°C) (Kibbe 2007). The fact that the saturated bound fraction is less than 1 could be due to (1) a common feature of filter-binding assays in which the process of physical separation disrupts bound species or (2) a stable population of protein-free crRNA:DNA complexes. In any case, the important conclusion to be drawn from these data is that each protospacer exhibits the same fraction bound regardless of the presence of mismatches at positions 19 and 20 in the crRNA. Thus, the crRNA-dependent effects seen in **Fig. 3A** and **Supp. Fig. 3B** must emerge from fundamental differences in conformational dynamics and not from differences in binding occupancy of Cas12a/crRNA on the DNA probe.

**Supplementary Figure 3B. Effect of R-loop truncation on permanganate reactivity of the A/T tract.** Permanganate reactivity of the A/T tract in a 20-nt R-loop and an 18-nt R-loop. In **Fig. 3A**, protospacer 3 is annotated as “DNA substrate 1;” protospacer 4 is annotated as “DNA substrate 2;” and crRNA 3 is the depicted crRNA. Permanganate experiments were conducted as in **Fig. 2B** (2 minutes, 30°C). See **Supp. Note 1** for description of the parameters plotted on the y-axis. Columns and associated error bars indicate the mean and standard deviation of three replicates. Columns 1, 2, 4, and 6 are equivalent to the data shown in **Fig. 3A**. Columns 3 and 5 use a crRNA with compensatory mutations at positions 19-20, showing that the effect is dependent upon base pairing topology and not a particular sequence.

**Supplementary Figure 3C. Effect of PAM-distal mismatches on non-target-strand and target-strand cleavage kinetics and position with fully duplex DNA targets.** 100 nM AsCas12a and 120 nM crRNA were incubated with 1 nM of DNA target at 37°C for 20 seconds, 1 minute, 5 minutes, and 30 minutes, prior to quenching and resolution by denaturing PAGE. Each group of four lanes corresponds to a different DNA target, with varying numbers of PAM-distal mismatches with respect to the crRNA. Indicated above each group of four lanes is the number of base pairs of complementarity between the TS and the crRNA spacer, starting with the base immediately adjacent to the PAM. All DNA targets in this gel were fully duplex (not pre-unwound/bubbled), resulting in enhanced discrimination against PAM-distal mismatches as compared to the bubbled DNA targets in **Fig. 3B** and **Supp. Fig. 3D**.

**Supplementary Figure 3D. Effect of PAM-distal mismatches on non-target-strand and target-strand cleavage kinetics and position with bubbled DNA targets.** 100 nM AsCas12a and 120 nM crRNA were incubated with 1 nM of DNA target at 37°C for 0 seconds, 15 seconds, 2 minutes, 10 minutes, and 1 hour, prior to quenching and resolution by denaturing PAGE. Each time series corresponds to a different DNA target, bearing varying numbers of PAM-distal mismatches with respect to the crRNA. Indicated above each time series is the number of base pairs of complementarity between the TS and the crRNA spacer, starting with the base immediately adjacent to the PAM. For the time series lacking an asterisk, the DNA target was fully duplex (as in **Supp. Fig. 3C**). For the time series that bear asterisks, the DNA target contained a bubble across the region of crRNA:TS complementarity (as illustrated in **Fig. 3B**), which stabilized the R-loop. In the top panel, the NTS was 5'-radiolabeled. In the bottom panel, the TS was 5'-radiolabeled.

**Supplementary Figure 3E. Determinants of altered target-strand cleavage kinetics and position.** 100 nM AsCas12a and 120 nM crRNA were incubated with 1 nM duplex DNA target radiolabeled on the 5' end of the target strand at 37°C for 0 s, 15 s, 1 m, 4 m, 15 m, or 1 h, prior to quenching and resolution by denaturing PAGE. The 20-nt target sequence immediately adjacent to the PAM is shown below the crRNA spacer sequence used in each experiment. Red letters indicate TS:crRNA mismatches. Green letters indicate compensatory changes in the crRNA to restore a 20-nt match. The final timepoint of each reaction is reproduced in the gel on the right, for side-by-side comparison of the cleavage site distributions. "Fraction cleaved" is defined as (sum of the volume of all bands below the uncleaved band)/(total volume in lane). Data were fit to an exponential decay ($y=(y_0\text{-plateau})*\exp(-k*x)+\text{plateau}$), with $y_0$ constrained to 0 and the plateau value constrained to ≤1. A representative replicate is shown. The value of $k_{obs}$ for each time course is as follows: **A** (0.12 $s^{-1}$), **B** (0.0093 $s^{-1}$), **C** (0.0094 $s^{-1}$), **D** (0.16 $s^{-1}$), **E** (0.015 $s^{-1}$). The precise value of $k_{obs}$ for **A** and **E** should be interpreted with caution due to poor sampling of informative timepoints.

**Supplementary Figure 3F. Non-target-strand cut-site distribution with a shrinking R-loop.** Final timepoint (1 hour) of each time series in the non-target-strand gel shown in the top panel of **Supp. Fig. 3D**, shown side-by-side for visual comparison—analogous to the final timepoints for the target strand shown in **Fig. 3B**.

**Supplementary Figure 3G. Kinetics of target-strand cleavage in DNA targets with various sequences in the R-loop flank.** Experiment performed as described in legend to **Fig. 3C**. 100 nM AsCas12a and 120 nM crRNA were incubated with 1 nM of DNA target at 25°C for 0 seconds, 15 seconds, 30 seconds, 1 minute, 2 minutes, 4 minutes, or 10 minutes, prior to quenching and resolution by denaturing PAGE. All DNA targets were 5'-radiolabeled on the TS. The NTS was pre-gapped from positions 14-18 but complementary to the TS at positions 1-13 and 19-20. In each lane, the DNA target was varied to contain different sequences in the RLF, which either formed a perfect duplex (substrates A, C, and E) or contained a 3-bp NTS:TS mismatch (substrates B, D, and F). "Fraction cleaved" is defined as (sum of the volume of all bands below the uncleaved band)/(total volume in lane). Data were fit to an exponential decay ($y=(y_0\text{-plateau})*\exp(-k*x)+\text{plateau}$), with $y_0$ constrained to 0. A representative replicate (n=3) is shown. The value of $k_{obs}$ ± SD for each time course is as follows: **A** [0.092 ± 0.012 $s^{-1}$], **B** [0.145 ± 0.007 $s^{-1}$], **C** [0.0059 ± 0.0006 $s^{-1}$], **D** [0.137 ± 0.002 $s^{-1}$], **E** [0.024 ± 0.002 $s^{-1}$], **F** [0.061 ± 0.013 $s^{-1}$]. The rate constants for **B** and **D** should be interpreted with caution due to poor sampling of informative timepoints.

**Supplementary Figure 4A. Permanganate reactivity of the A/T tract in R-loops formed by dCas12a or dCas9.** Permanganate experiments were conducted as in **Fig. 2B** (2 minutes, 30°C). “Pre-gapped” indicates the presence of a 5-nt gap in the non-target strand (see **Appendix B**) (the NTS gap in the dCas9 target is unrelated to the cut that would normally be formed by a nuclease-active Cas9—instead, it was designed to be analogous to the NTS gap formed by AsCas12a in an R-loop of the opposite topology, at positions 14-18). “Pre-unwound” indicates the presence of a stretch of NTS:TS mismatches in the DNA substrate (20-nt bubble throughout region of RNA complementarity); asterisks highlight the constructs that contain these NTS:TS mismatches. The sequence of protospacer 5 is identical to that of protospacer 2 except for a change in the PAM, which is not expected to affect conformational dynamics at the A/T tract (besides in permitting dCas9 binding). See **Supp. Note 1** for description of the parameters plotted on the y-axis. Columns and associated error bars indicate the mean and standard deviation of three replicates. Experiments **A**, **B**, **E**, **F**, **G**, and **J** are equivalent to the data shown in **Fig. 4A**.

**Supplementary Figure 4B. dCas9 binds tightly to pre-gapped DNA targets.** The affinity of dSpCas12a/crRNA for a cognate DNA target was assessed by an electrophoretic mobility shift assay (EMSA) and a filter-binding (FB) assay. dSpCas9 was titrated in a solution with fixed [sgRNA] (750 nM) and [DNA probe] (100 pM), followed by separation of protein-bound DNA from free DNA. The EMSA indicated that the binding conditions yield one major protein-bound species. “Fraction bound” is defined as (background-subtracted volume of upper band)/(total background-subtracted lane volume) for the EMSA and (background-subtracted volume of Protran spot)/(total background-subtracted volume of Protran spot + Hybond N+ spot) for the filter-binding assay. The value of “fraction bound” was 0 at [dSpCas9]=0 for both substrates and both assays (not shown on plot due to the logarithmic x-axis). All data shown are from one representative replicate (n=3). Protospacer 6 is identical to protospacer 1 (used for AsCas12a), except the PAM has been substituted with a SpCas9 PAM. Protospacer 5 is a modified version of protospacer 6, with differences highlighted in red (equivalent to the protospacer-1-to-protospacer-2 modifications). The “intact” protospacer is as shown in the sequence schematic. The “pre-gapped” protospacer is missing nt 14-18 of the NTS (as measured from the PAM, design). The NTS gap is unrelated to the cut that would normally be formed by a nuclease-active Cas9—instead, it was designed to be analogous to the NTS gap formed by AsCas12a in an R-loop of the opposite topology, at positions 14-18. However, this DNA substrate still binds tightly to dSpCas9. Thus, the failure of dSpCas9 to significantly distort the RLF in **Fig. 4A** is due to a fundamental difference in conformational dynamics and not to a failure to bind.

**Supplementary Figure 4C. Permanganate reactivity of the A/T tract in protein-free R-loops of various sequences.** Permanganate experiments were conducted as in **Fig. 2B** (2 minutes, 30°C). See **Supp. Note 1** for description of the parameters plotted on the y-axis. Columns and associated error bars indicate the mean and standard deviation of three replicates. In all schematics, RNA molecules are outlined in orange, and DNA molecules are outlined in black. Circled "P" indicates a 5′-phosphate. All sequences, when read right-side up, go from 5′ on the left to 3′ on the right. The terms "Cas12a-like" and "Cas9-like" are descriptors only of each substrate's R-loop topology (the end of the RNA next to the boundary of interest is a 3′ end or a 5′ end, respectively)—both kinds of substrates contain a Cas12a PAM and a Cas12a-like NTS gap. These results imply that the asymmetry in RLF stability is a fundamental feature of R-loop structure and not a peculiarity of the original tested sequence.

**Supplementary Figure 4D. Effect of RNA end chemistry on permanganate reactivity of the A/T tract in protein-free R-loops.** Permanganate experiments were conducted as in **Fig. 2B** (2 minutes, 30°C), varying only the nature of the RNA molecule added to the pre-gapped/pre-unwound DNA substrate. See **Supp. Note 1** for description of the parameters plotted on the y-axis. Columns and associated error bars indicate the mean and standard deviation of three replicates. In both schematics, RNA molecules are outlined in orange, and DNA molecules are outlined in black. Circled "P" indicates a 5′-phosphate. All sequences, when read right-side up, go from 5′ on the left to 3′ on the right. "OH" indicates a hydroxyl. "Phos." indicates a phosphate. "Cyc. phos." indicates a 2′/3′-cyclic phosphate. "IVT/rz" indicates that the RNA oligo was synthesized in an enzymatic *in vitro* transcription reaction, with ribozymes on both ends that cleaved to yield homogeneous termini. "Chem." indicates that the RNA oligo was chemically synthesized by a commercial source. The terms "Cas12a-like" and "Cas9-like" are descriptors only of each substrate's R-loop topology (the end of the RNA next to the boundary of interest is a 3′ end or a 5′ end, respectively)—both kinds of substrates contain a Cas12a PAM and a Cas12a-like NTS gap.

**Supplementary Figure 4E. Asymmetry in R-loop flank stability is also a feature of intact R-loops.** Permanganate experiments were conducted as in **Fig. 2B** (2 minutes, 30°C). See **Supp. Note 1** for description of the parameters plotted on the y-axis. Columns and associated error bars indicate the mean and standard deviation of three replicates. In all schematics, RNA molecules are outlined in orange, and DNA molecules are outlined in black. All sequences, when read right-side up, go from 5′ on the left to 3′ on the right. These experiments provide the strongest point of comparison between the Cas12a-like and Cas9-like R-loop architecture, as the only component varied across conditions is which strand of an identical DNA bubble is hybridized to RNA (compare the cleaved R-loops, in which the position of the gap must be moved to the opposite strand, yielding slightly different baseline permanganate reactivity).

**Supplementary Figure 4F. Effect of overhanging non-target-strand nucleotides on permanganate reactivity of the A/T tract in protein-free R-loops.** Permanganate experiments were conducted as in **Fig. 2B** (2 minutes, 30°C). See **Supp. Note 1** for description of the parameters plotted on the y-axis. Columns and associated error bars indicate the mean and standard deviation of three replicates. In all schematics, RNA molecules are outlined in orange, and DNA molecules are outlined in black. Circled "P" indicates a 5′-phosphate. All sequences, when read right-side up, go from 5′ on the left to 3′ on the right. These experiments probe the role of the 2-nt NTS overhang (immediately adjacent to the R-loop flank) in distortion of the A/T tract. When present, this dinucleotide results in a "flapped" R-loop flank (RLF) terminus, and when absent, the RLF terminus is "flush." The terms "(Cas)12a-like" and "(Cas)9-like" are descriptors only of each substrate's R-loop topology (the end of the RNA next to the boundary of interest is a 3′ end or a 5′ end, respectively)—both kinds of substrates contain a Cas12a PAM and a Cas12a-like NTS gap. These results show that the presence of the overhang can affect the absolute magnitude of distortion, but the nature of the asymmetry is unaffected.

**Supplementary Figure 4G. 2-aminopurine fluorescence measurements confirm asymmetry in conformational dynamics of R-loop flanks.** Columns and error bars show mean and standard deviation of three replicates. In all schematics, RNA molecules are outlined in orange, and DNA molecules are outlined in black. Circled “P” indicates a 5′-phosphate. All sequences, when read right-side up, go from 5′ on the left to 3′ on the right. The terms “Cas12a-like” and “Cas9-like” are descriptors only of each substrate’s R-loop topology (the end of the RNA next to the boundary of interest is a 3′ end or a 5′ end, respectively)—both kinds of substrates contain a Cas12a PAM and a Cas12a-like NTS gap. The absolute values of 2-AP fluorescence intensity have a wide range across different DNA probes, likely due to local sequence context or, in the case of the ssDNA control in the bottom right panel, perhaps the stable population of a conformation that enhances 2-AP fluorescence. Given this variation, it is important to use the perfect DNA duplex (**C**) and the pre-gapped/pre-unwound DNA bubble (**D**) conditions as benchmarks—on the continuum from **C** to **D**, where does **E** lie? For condition **E** in the Cas12a-like topologies, the 2-AP fluoresces as if the RNA were absent. For condition **E** in the Cas9-like topologies, 2-AP fluorescence is quenched and approaches the intensity of the fully duplex control.

**Supplementary Figure 5A. Thermal stability determination for nicked dumbbell substrates and their constituent hairpins.** Data from representative replicates of refolding experiments (small black dots) are overlaid on a best-fit curve (thick blue line) comprising a Boltzmann sigmoid with inclined baselines (y=(m*x+b)+((n*x+c)-(m*x+b))/(1+exp(($T_m$-x)/slope))). Because the dumbbells contain two separate duplexes that can, in principle, fold and unfold independently of each other, each of these molecules likely has more than two states. Thus, while there is no obvious visual sign of multiple transitions in the refolding curves, we did not attempt to extract thermodynamic parameters from the slope, and the $T_m$ (written in the center of each plot) should only be used as a point of comparison rather than a determinant of a defined conformational ensemble. While the two RNA:DNA hairpins have slightly different $T_m$ values, the much larger discrepancy in $T_m$ of the nicked dumbbells is probably mostly due to the nature of duplex juxtaposition.

**Supplementary Figure 5B. Molecular dynamics simulations of the Cas12a-like and Cas9-like interhelical junctures, Sequence 2.** Schematics and data are presented as described in the legend to **Figure 5B**, which depicted simulation of Sequence 1. For Sequence 2, the nucleobases probed at the 3' R-loop boundary frequently exhibited unstacking or poorly stacked conformations (although to a lesser extent than in Sequence 1), while those probed at the 5' R-loop boundary were stably stacked over the course of the simulation. These results suggest that the difference in stacking instability detected for the interhelical junctures (with Sequence 1 or Sequence 2) are due to the difference in juncture topology rather than the identities of the monitored nucleobases (pyrimidine/pyrimidine vs. purine/purine).

**Supplementary Figure 5C. Effect of crRNA spacer mimic modifications on permanganate reactivity of the A/T tract in protein-free R-loops.** Permanganate experiments were conducted as in **Fig. 2B** (2 minutes, 30°C). See **Supp. Note 1** for description of the parameters plotted on the y-axis. Columns and associated error bars indicate the mean and standard deviation of three replicates. In all sequence schematics, RNA molecules are outlined in orange, and DNA molecules are outlined in black. Circled "P" indicates a phosphate. All sequences, when read right-side up, go from 5′ on the left to 3′ on the right. The series of rectangles shown below each column represent a variant of the crRNA spacer mimic added to the R-loop depicted on the left. When uridine ribonucleotides where changed to 2′-deoxyribonucleotides, the nucleobase was also changed to thymine (to minimize cost of synthesis). The terms "Cas12a-like" and "Cas9-like" are descriptors only of each substrate's R-loop topology (the end of the RNA next to the boundary of interest is a 3′ end or a 5′ end, respectively)—both kinds of substrates contain a Cas12a PAM and a Cas12a-like NTS gap.

**Supplementary Figure 7A. Substrate specificity of Cas12a *trans*-active holoenzyme and S1 nuclease.** *Top panel*: Enzymes were first titrated in an activity assay with a radiolabeled ssDNA substrate to determine what concentration to use in substrate specificity assays. A ssDNA oligo (the same one shown in the other panels) was 5'-radiolabeled and incubated with varying concentrations of either AsCas12a/crRNA/pre-cleaved DNA activator (all components held equimolar at the indicated concentration) or S1 nuclease for 30 minutes at 30°C, followed by quenching. Products were resolved by denaturing PAGE and quantified from the phosphorimage. "U" on the x-axis of S1 nuclease refers to the units defined by the enzyme manufacturer. To achieve 90% cleavage of the ssDNA substrate in the given time course, 115 nM AsCas12a holoenzyme or 0.513 U/µL S1 nuclease was required. *Middle panels*: Phosphorimage of AsCas12a or S1 nuclease cleavage products, resolved by denaturing PAGE. *Trans*-active AsCas12a holoenzyme (115 nM of each component: protein, crRNA, pre-cleaved activator) or S1 nuclease (0.513 U/µL) was incubated with 1 nM of the indicated substrate for 2 hours at 30°C prior to quenching. Substrate (a) was a single-stranded DNA oligonucleotide with no homology to the crRNA. To generate substrates (b) through (l), substrate (a) was hybridized to a variety of unlabeled complementary DNA oligonucleotides. Substrate (c) contained a nick. Substrates (d), (e), and (f) contained gaps of 1, 4, and 8 nucleotides (nt), respectively. Substrates (g), (h), and (i) contained bubbles of 1, 4, and 8 nt, respectively. Substrates (j), (k), and (l) contained bulges of 1, 4, and 8 nt, respectively. *Bottom panel*: Quantifications of cleavage from phosphorimages. "Fraction cleaved" is defined as (sum of the volume of all bands below the uncleaved band)/(total volume in lane).

**Supplementary Figure 7B. Denaturing PAGE analysis of AsCas12a cleavage products.** *Top panel*: The leftmost lane in each gel is a 1-hour timepoint with AsCas12a D908A (mutation in the RuvC active site). The timepoints shown for the reactions with WT enzyme are 0 s, 5 s, 10 s, 20 s, 40 s, 80 s, 10 m, 1 h. The ladder was made by digestion of the relevant substrate with a cocktail of non-specific DNases (see Methods). The rightmost lane is the same probe cleaved by the site-specific restriction enzyme TseI. *Bottom panel*: The same products run on a gel for a shorter amount of time to demonstrate the absence of faster-migrating species. This gel lacks the ladder samples.

**Supplementary Figure 7C. Steady-state kinetic analysis of AsCas12a *trans* DNase activity, as measured by fluorophore dequenching.** Raw data of a representative replicate are shown in the middle panel. The $V_0$ values of the representative replicate are plotted in the bottom panel, along with a hyperbolic fit ($y=v_{max}*x/(K_M+x)$). The table shows the average and standard deviation of kinetic parameters, determined from three independent replicates. The $k_{cat}$ value reported in the table is based on an upper limit of 10 nM *trans*-active AsCas12a holoenzyme. See the figure legend to **Supp. Fig. 7D** for a comparison of *trans* and *cis* cleavage kinetics.

**Supplementary Figure 7D. Enzyme-concentration dependence of AsCas12a *cis* DNA cleavage kinetics.** 250 pM of a *cis* duplex DNA target, radiolabeled on the 5' end of the NTS, was incubated with 240 nM crRNA and various concentrations of AsCas12a at 37°C for the following timepoints: 0 s, 5 s, 10 s, 20 s, 40 s, 10 m. “Fraction cleaved” is defined as (sum of the volume of all bands below the uncleaved band)/(total volume in lane). Data were fit to an exponential decay ($y=(y_0\text{-plateau})*\exp(-k*x)+\text{plateau}$), with $y_0$ constrained to 0. At the lower values of [AsCas12a], an early plateau indicates an underestimation of the true enzyme concentration, due to loss of enzyme across extensive serial dilution. Despite this experimental deficiency, these data show that enzyme-substrate association is not rate-limiting when [AsCas12a] is 100 nM—i.e., when [AsCas12a] is 100 nM, association can be approximated as instantaneous, and cleavage rate constants reflect unimolecular processes that follow binding. Based on these experiments, initial *cis* cleavage events occur at ~0.12 $s^{-1}$ at the reagent concentrations used for cleavage mapping experiments. Considering the steady-state kinetic measurements in **Supp. Fig. 7C**, an upper limit on *trans* cleavage rate can be calculated as $(k_{cat}/K_M)*[E]$, where [E] has an approximate upper limit of 10 nM (for radiolabeled-NTS experiments, in which cold TS is present at 10 nM, and assuming ssDNA-activated complexes have similar catalytic efficiency to dsDNA-activated complexes), yielding a rate of ~0.002 $s^{-1}$ (>50-fold less than the *cis* cleavage rate). This *trans* cleavage rate is an upper limit because radiolabeled DNA is mostly duplex and likely protected by the protein in non-duplex regions (Swarts et al. 2017). Thus, *trans* cleavage probably only occurs to a detectable extent on PAM-distal DNA fragments that are released from the enzyme following *cis* cleavage.

**Supplementary Figure 7E. Concentration dependence of various modes of DNase activity.** In biochemical reactions containing DNA, crRNA, and AsCas12a, a cut in a DNA molecule could be attributed to one of three distinct modes of RuvC DNase activity: *cis* cleavage of an enzyme's own bound R-loop; *trans* cleavage (by an R-loop-activated complex) of free DNA or DNA in a different complex; or *trans* cleavage by an excess of DNA-free Cas12a/crRNA complex (which is shown in this figure to be catalytically competent, albeit inefficient). Various concentrations of crRNA, AsCas12a, and DNA activator were incubated with 2 nM of a radiolabeled single-stranded DNA oligonucleotide for 1 hour at 37°C prior to quenching and resolution by denaturing PAGE. "Fraction cleaved" is defined as (sum of the volume of all bands below the uncleaved band)/(total volume in lane). We show here that both DNA-activated and DNA-free modes of *trans* cleavage are dependent upon enzyme concentration in the nanomolar range—i.e., unlike *cis* cleavage, which occurs with saturated binding kinetics at the concentrations used (see **Supp. Fig. 7D**), *trans* cleavage remains concentration-dependent in this range. Therefore, DNase cleavage products that appear with kinetics that are independent of enzyme concentration can be uniquely attributed to PAM-dependent *cis* cleavage (see **Supp. Fig. 7F**).

**Supplementary Figure 7F. Concentration dependence of phosphodiester-mapped cleavage events.** Each graph follows the kinetics of appearance/disappearance of a cleavage product at the indicated phosphodiester. “Fraction cleaved” is defined as (band volume for a given cleavage product)/(total volume in lane). For example, “NTS 5', 11/12” displays the fraction of 5'-radiolabeled NTS that was detected to have been cleaved at the 11/12 dinucleotide for each timepoint. Blue circles are equivalent to the data shown in **Fig. 7**, with error bars denoting standard deviation across three replicates. Red squares and green triangles indicate variants of this experiment in which the total concentration of either DNA-bound holoenzyme (red squares) or AsCas12a/crRNA (green triangles) was decreased. The fact that almost all species exhibit equivalent kinetics of appearance/disappearance in all three conditions (with the potential exception of the phosphodiesters on the portion of the PAM-distal NTS fragment that remains single-stranded after release by the enzyme, which exhibit slight concentration-dependence on longer timescales) indicates that we are observing mostly *cis* cleavage events, as explained in **Supp. Fig. 7E**.

**Supplementary Figure 7G. Cleavage product mapping for FnCas12a.** The evolution of the pattern of *cis* cleavage over time for FnCas12a was assayed analogously to AsCas12a, as in **Fig. 7**. The pattern of cleavage is largely similar to that of AsCas12a. However, the NTS gap of FnCas12a widens more gradually than that of AsCas12a, and the TS cleavage distribution is more tightly constrained to a single phosphodiester. No DNA cleavage was detected on either strand when a RuvC-inactivated FnCas12a mutant (D917A) was used.

**Supplementary Figure 8A. Non-target-strand cleavage precedes target-strand cleavage for AsCas12a and Cas12i1.** 100 nM AsCas12a or Cas12i1 was incubated with 120 nM cognate crRNA and 2 nM radiolabeled duplex DNA target for 1 hour at 37°C, followed by quenching, denaturing PAGE, and phosphorimaging. For AsCas12a, the reaction was conducted in 5 mM $CaCl_2$. For Cas12i1, the reaction was conducted in 5 mM $MgCl_2$. In the duplex diagrams, red shading indicates the presence of a phosphorothioate (PS) tract across the standard cleavage sites. Blue indicates phosphodiester (PO) linkages within the TS. Gray indicates phosphodiester linkages within the NTS. For both AsCas12a and Cas12i1, the nature of the linkages in the TS has no apparent effect on NTS cleavage. However, the presence of phosphorothioates in the NTS inhibits cleavage of both the NTS and the TS. Trace TS cleavage is observed for Cas12i1 in the PS-NTS condition (lane 6)—it is unclear whether this is due to TS cleavage prior to NTS cleavage or to incomplete duplex formation, as the trace cleavage event is shifted to the site cleaved during ssDNA-targeting (lane 7).

**Supplementary Figure 8B. Cleavage at phosphorothioates can be selectively slowed by substitution of $CaCl_2$ for $MgCl_2$.** 100 nM AsCas12a and 120 nM crRNA were incubated with 1 nM radiolabeled duplex DNA target at 37°C, followed by quenching (at timepoints 0, 5 s, 15 s, 1 m, 5 m, 10 m, 30 m, 1 h) and resolution by denaturing PAGE. Substrate diagrams are colored as in **Supp. Fig. 8A**. The top panel shows the experiment done in cleavage buffer with 5 mM $MgCl_2$. The bottom panel shows the experiment done in cleavage buffer with 5 mM $CaCl_2$—at the end of each time course on this gel, the 1-hour timepoint of the $MgCl_2$ experiment is included for visual comparison. “Fraction cleaved” is defined as (sum of the volume of all bands below the uncleaved band)/(total volume in lane). Data were fit to an exponential decay ($y=(y_0\text{-plateau})*\exp(-k*x)+\text{plateau}$), with $y_0$ constrained to 0. The plateau value was constrained to 1 for those time courses that did not exceed fraction cleaved = 0.5 by the 1-hour timepoint. Rate constants (with 95% confidence intervals) are shown in the table below the gels. It is unclear why cleavage of a phosphorothioated TS occurs more rapidly than cleavage of a phosphorothioated NTS, although it is conceivable that this is an intrinsic feature of the enzyme cleavage pathway when the chemical transformation is rate-limiting. Considering only effects on the NTS, the calcium substitution decreases the phosphodiester cleavage rate by a factor of 57 and decreases the phosphorothioate cleavage rate by a factor of 730, resulting in a 13-fold increase in selectivity for phosphodiesters over phosphorothioates and yielding kinetics slow enough to resolve by manual pipetting.

**Supplementary Figure 8C. Interference complexes are stable in the presence of $CaCl_2$ and with a phosphorothioated DNA target.** Using a filter-binding assay, we assessed the affinity of dAsCas12a/crRNA for cognate DNA targets, either fully phosphodiester (PO) or containing phosphorothioate (PS) linkages across the NTS and TS cleavage sites, in the presence of either 5 mM $MgCl_2$ or 5 mM $CaCl_2$. Substrate diagrams are colored as in **Supp. Fig. 8A**. "Fraction bound" is defined as (background-subtracted volume of Protran spot)/(total background-subtracted volume of Protran spot + Hybond N+ spot). The value of "fraction bound" was 0 at [dAsCas12a]=0 for both substrates and both assays (not shown due to the logarithmic x-axis). All data shown are from a representative replicate (n=3). When appropriate, data were fit to the sum of a hyperbola and a line ($y=B_{max}*x/(K_D+x)+NS*x$), where NS describes a non-specific binding mode. It is common to see $B_{max}$ values below 1 in filter binding assays, in which the process of physical separation can disrupt bound species. $K_D$ for the PO substrate in $MgCl_2$ was 8.1 nM ± 0.8 (SD) (n=3). Data from the other binding conditions indicated that the $K_D$ was near or below [DNA probe], preventing accurate $K_D$ determination by hyperbolic fitting. For unknown reasons, the $Mg^{2+}$→$Ca^{2+}$ and the PO→PS substitutions stabilized the ribonucleoprotein:DNA interaction both separately and together.

**Supplementary Figure 8D. Non-target-strand variants used in gap-dependence experiments.** Nucleotides 7-22 of the NTS are shown for each variant. A red circle indicates a phosphorothioate linkage. A gray circle indicates a phosphodiester linkage. A 5′ phosphate was placed on all PAM-distal NTS fragments to recapitulate the end chemistry left by the RuvC-catalyzed hydrolysis reaction.

**Supplementary Figure 8E. Phosphorimage and quantification of non-target-strand gap-dependence experiments, in $CaCl_2$.** Experiment was performed as described in legend to **Fig. 8A**. Substrates indicated by a letter are as schematized in **Supp. Fig. 8D**. When a given category has more than one substrate (e.g., 3-nt gap includes substrates M, N, O, and P), the first listed substrate (M) is shown as a blue bar, the second (N) as a red bar, the third (O) as a green bar, and the fourth (P) as a pink bar. “Fraction cleaved” is defined as (sum of the volume of all bands below the uncleaved band)/(total volume in lane).

**Supplementary Figure 8F. Phosphorimage and quantification of non-target-strand gap-dependence experiments, in $MgCl_2$.** Experiment was performed as described in legend to **Fig. 8A**, except 5 mM $MgCl_2$ was used instead of $CaCl_2$, and the reaction was quenched after only 10 seconds at 37°C. Substrates indicated by a letter are as schematized in **Supp. Fig. 8D**. When a given category has more than one substrate (e.g., 3-nt gap includes substrates M, N, O, and P), the first listed substrate (M) is shown as a blue bar, the second (N) as a red bar, the third (O) as a green bar, and the fourth (P) as a pink bar. “Fraction cleaved” is defined as (sum of the volume of all bands below the uncleaved band)/(total volume in lane).

**Supplementary Figure 8G. Phosphorimage and quantification of non-target-strand gap-dependence experiments, in $MgCl_2$, with radiolabeled *trans* substrate.** Extent of *trans* cleavage by wild type AsCas12a in the presence of various DNA activator variants, as resolved by denaturing PAGE. Cas12a ternary complex (final concentrations: 100 nM AsCas12a, 120 nM crRNA, pre-hybridized 120 nM TS / 240 nM NTS) was assembled with each of the indicated NTS variants and combined with 2 nM (final concentration) radiolabeled non-specific *trans* ssDNA target in cleavage buffer (5 mM $MgCl_2$). These reactions were then incubated for 30 minutes at 37°C prior to quenching and resolution by denaturing PAGE. Control lanes on the left contain some combination of intact NTS/TS with phosphodiesters (PO) or phosphorothioates (PS) across the standard cleavage sites; "pc" stands for pre-cleaved (only PAM-proximal cleavage products: NTS truncated after nt 13, TS truncated after nt 22). Reactions without NTS contained 120 nM of a non-specific DNA oligonucleotide to account for substrate competition. All lanes indicated by a letter (C-Y) contained an NTS variant (see **Supp. Fig. 8D**) along with a TS containing phosphorothioates across the standard cleavage sites. When a given category has more than one substrate (e.g., 3-nt gap includes substrates M, N, O, and P), the first listed substrate (M) is shown as a blue bar, the second (N) as a red bar, the third (O) as a green bar, and the fourth (P) as a pink bar. "Fraction cleaved" is defined as (sum of the volume of all bands below the uncleaved band)/(total volume in lane).

**Supplementary Figure 8H. Phosphorimages and quantification of pre-gapped non-target-strand experiments.** Experiment was performed as described in legend to **Fig. 8B**. “Fraction cleaved” is defined as (sum of the volume of all bands below the uncleaved band)/(total volume in lane). Data were fit to an exponential decay ($y=(y_0-plateau)*exp(-k*x)+plateau$), with $y_0$ constrained to 0. The plateau value was constrained to 1 for those time courses that did not exceed fraction cleaved = 0.5 by the 1-hour timepoint. The exponential decay constant k is reported as $k_{obs}$ in **Fig. 8B**. The data shown here are from a representative replicate (n=3).

**Supplementary Figure 8I. Affinity measurements for RNA-guided interaction of AsCas12a mutants with dsDNA.** The affinity of AsCas12a/crRNA for a cognate DNA target was assessed by a filter-binding assay, with three variants of AsCas12a. AsCas12a was titrated in a solution with fixed [crRNA] (750 nM) and [DNA probe] (100 pM), in binding buffer containing $MgCl_2$, followed by separation of protein-bound DNA from free DNA on membranes. “Fraction bound” is defined as (volume of Protran spot)/(total volume of Protran spot + Hybond N+ spot). The value of “fraction bound” was 0 at [AsCas12a]=0 for all protein variants (not shown due to the logarithmic x-axis). All data shown are from a representative replicate (n=3). Data were fit to the sum of a hyperbola and a line ($y=B_{max}*x/(K_D+x)+NS*x$), where NS describes a non-specific binding mode. It is common to see $B_{max}$ values below 1 in filter binding assays, in which the process of physical separation can disrupt bound species. $K_D$ for AsCas12a D908A was 11 nM ± 3 (SD) (n=3). $K_D$ for AsCas12a R1226A was 2.7 nM ± 0.2 (SD) (n=3). $K_D$ for AsCas12a WT was 1.86 nM ± 0.04 (SD) (n=3).

**Supplementary Figure 8J. Non-target-strand cleavage product mapping for AsCas12a R1226A.** The evolution of the pattern of *cis* cleavage over time for AsCas12a R1226A was assayed analogously to WT AsCas12a, as in **Fig. 7**. At the 1-hour timepoint, AsCas12a R1226A had made at least one cut in ~70% of the assayed NTS molecules, which matches the average fraction cleaved by WT AsCas12a at the 5-second timepoint. The cleavage site distributions in each of these experiments can be compared by calculating the probability density of each cleavage product as: (volume of band corresponding to cleavage at phosphodiester X/Y)/(total volume of all cleaved products), such that the area under each distribution is 1. Unlike the WT distribution, for which the 5′-mapped and 3′-mapped distributions are mostly non-overlapping, the R1226A distributions have significant overlap at dinucleotides 14/15, 15/16, and 16/17. In our ensemble biochemical experiment, this overlap is consistent with (although not uniquely explained by) the presence of a large population of molecules that have been cleaved exactly once (i.e., have not undergone gap formation). If this explanation is correct, these data point to an accumulation of AsCas12a R1226a complexes with a once-cut NTS, supporting the model that gap formation is disproportionately slow (and perhaps rate-limiting for TS cleavage) for this mutant.

**Supplementary Figure 8K. Cleavage product mapping for Cas12i1.** The evolution of the pattern of *cis* cleavage over time for Cas12i1 was assayed analogously to WT AsCas12a, as in **Fig. 7**. The pattern of cleavage is largely similar to that of AsCas12a, although cleavage kinetics are dramatically slower, and disproportionately so for TS cleavage. The NTS gap of Cas12i1 is wider (8 nt) than that of AsCas12a (5 nt) at 1 hour. No DNA cleavage was detected on either strand when a RuvC-inactivated Cas12i1 mutant (D647A) was used.
