## Supplementary material for "CRISPR-Cas12a exploits R-loop asymmetry to form double-strand breaks": Supp_Note_1.pdf

**Supplementary Note 1. Quantification of permanganate reactivity and limitations of interpretation.**

In this work, data describing permanganate reactivity are presented in three ways:

1. Raw phosphorimages of denaturing PAGE analysis of DNA substrates treated with permanganate and piperidine.
2. "Permanganate reactivity index" (PRI) of individual thymine nucleobases. This metric is determined from the raw phosphorimages. It is an approximation of the absolute rate of oxidation at a given thymine, linearly normalized such that PRI = 1 describes a thymine that is fully single-stranded. Thus, a thymine with PRI = 0.4 is estimated to have been oxidized twice as fast as a thymine with PRI = 0.2.
3. "Fraction oxidized in A/T tract" (FO). This metric is a mathematical transformation/combination of the PRI of all thymines in the R-loop flank of a given DNA substrate. It is an approximation of the total fraction of DNA molecules (the two strands of DNA forming the R-loop flank are referred to here as a single "molecule") that, at the moment of quenching, have been oxidized on at least one of the nine thymines within the R-loop flank.

While visually inspecting phosphorimages from permanganate experiments, note that there are occasionally faint bands corresponding to strand cleavage at cytosines (permanganate oxidizes cytosines, albeit much more slowly than thymines) and at purines (which occurs during hydroxide treatment in the 3′-radiolabeling protocol). Such bands constituted a trivial fraction of the total lane volume and did not meaningfully affect analysis.

Additionally, visual inspection of the raw phosphorimages can be informative but should be approached with caution because the absolute volume of a given band is meaningless without considering other bands that may have detracted from its signal. For example, a strongly oxidized thymine (thymine 1) may yield only a faint band if another strongly oxidized thymine (thymine 2) lies between thymine 1 and the radiolabeled terminus of the DNA oligonucleotide. If all thymine oxidation events are independent of each other (i.e., thymine 1 has the same oxidation probability irrespective of whether thymine 2, or any other thymine, has been oxidized or not), the oxidation probability of thymine 1 can be reconstructed by considering the thymine-1 band only as a subpopulation of the bands above it on the gel. In other words, out of all the DNA molecules on the gel for which oxidation of thymine 1 would have been observable (i.e., cleavage fragments *at or above* the thymine-1 fragment on the gel), what fraction of those molecules *were* in fact oxidized at thymine 1? In reality, clusters of thymines have been observed to mutually enhance oxidation probability (Nomura and Okamoto 2008), so perfect independence *cannot* be assumed. Thus, the parameters described below are imperfect measures of the true rate of oxidation at each thymine.

Beyond uncertainty in the measurement, it is also unknown to what extent the probing technique is changing the fundamental biophysical features of the DNA structures. Notably, thymine’s reaction with permanganate breaks the planarity of the nucleobase and, consequently, its capacity to stack normally. In an A/T-rich sequence like our R-loop flank (**Fig. 2B**), an oxidation event at thymine 1 could, in principle, begin a chain reaction of oxidation events as each adjacent thymine successively loses planarity and unstacks, exposing its neighbor to the oxidant. If such chain reactions occurred quickly as compared to the timescale of the assay (2 minutes), the distribution of band volumes would be skewed toward thymine 9. In reality, the band volume distributions are skewed sharply toward thymine 1 (**Supp. Fig. 2C**), suggesting that, on the assayed timescale, the majority of oxidation events do *not* lead to additional oxidation events. Still, the possibility of chain reactions should be kept in mind when interpreting the observed permanganate reactivity patterns, in which reactivity decreases with distance from the R-loop edge (**Fig. 2B**). While these patterns are consistent with fraying duplex termini, the apparent "depth" of the fraying events should be interpreted as an upper limit on what would occur in a substrate unexposed to permanganate.

Finally, the structural determinants of permanganate reactivity should be considered carefully when using these data to draw conclusions about DNA conformation. While high permanganate reactivity is often associated with "single-strandedness" or "lack of base pairing," the reaction is more precisely dependent upon the ability of a permanganate molecule to approach the C5=C6 bond of the thymine nucleobase. This approach could be facilitated by assumption of a non-B-form helical geometry, global melting of the DNA duplex, or "flipping" of a thymine out of the duplex without dramatically affecting the helical geometry (Bui et al. 2003). Furthermore, a thymine lying on a duplex terminus could, in principle, be approached and attacked while base paired, albeit from a restricted angle. This possibility is especially important to consider for thymine 1 of our A/T-rich R-loop flank (**Fig. 2B**). The reactivity of this thymine varies in RNA-free DNA bubble controls that have different bubble sequences (**Supp. Fig. 4C**), perhaps reflecting differences in the propensities of individual (unpaired) neighboring bases to stack on the duplex-terminal thymine. Finally, because thymines within the RNA:DNA hybrid of R-loop structures have two possible base pairing partners (DNA versus crRNA), the conformational ensemble at these positions is highly complex, and we did not attempt to draw any structural conclusions from their oxidation rates.

Given the aforementioned caveats, the PRI and FO metrics described below should be interpreted as estimates rather than accurate measurements of rate and extent of reaction. Additionally, permanganate reactivity data should be considered alongside the orthogonal techniques used in this work to assess the structure, energetics, and conformational dynamics of interhelical junctures.

**Definitions of permanganate reactivity index (PRI) and fraction oxidized (FO)**

Let $v_i$ denote the volume of band $i$ in a lane with $n$ total bands (band 1 is the shortest cleavage fragment, band $n$ is the topmost band corresponding to the starting/uncleaved DNA oligonucleotide). The probability of oxidation at thymine $i$ is defined as:

$$p_i = \frac{v_i}{\sum_{j=i}^{n} v_j}$$

Note that this relationship allows determination of $p_i$ even if the values of $p_{1 \leq x < i}$ are unavailable (e.g. if the shortest cleavage products have been run off the bottom of the gel). Assuming thymine oxidation occurs with a uniform probability across the time course of permanganate application (see exponential curve in **Supp. Fig. 2C**), the rate constant associated with oxidation probability $p_i$ is defined as:

$$k_i = \frac{\ln(\frac{1}{1 - p_i})}{t}$$

where $t$ is the time of quenching. We found that across experimental replicates there was systematic variation in $k$ (e.g. $k$ was universally smaller in replicate 2 than in replicate 1 for any given thymine), likely due to variability in the oxidation activity of each new preparation of the potassium permanganate solution. To allow comparison across replicates, we normalized all values of $k$ to that of a reference thymine ($k_{ref}$) whose conformational dynamics were not expected to be affected by R-loop formation or associated substrate variations (the thymine 10 nt from the end of the 5′-radiolabeled oligo, present in all DNA substrates tested). For every set of replicate experiments, which each involved a new preparation of potassium permanganate solution, we determined the average rate constant of the reference thymine across all substrates ($\bar{k}_{ref}$). The global average of $k_{ref}$ across all experiments and all replicates ($\mu_{ref}$) was taken to be the true value of $k_{ref}$. The corrected value of $k_i$ for each thymine was then taken to be:

$$k_{i,corr} = \frac{\mu_{ref}}{\bar{k}_{ref}} k_i$$

The permanganate reactivity index was then calculated as:

$$PRI_i = \frac{k_{i,corr}}{k_{ss,corr}}$$

where $k_{ss,corr}$ is the reference-corrected oxidation rate constant for a thymine unassociated with a stable base-pairing partner. The value of $k_{ss,corr}$ used in our calculations was 0.79 min$^{-1}$, empirically determined for an arbitrarily chosen thymine within a DNA bubble (**Supp. Fig. 2C**). The estimated fraction of DNA molecules oxidized on at least one thymine within the R-loop flank (correcting to the value expected if the potassium permanganate solution had its average oxidation activity) was then calculated as:

$$p_{i,corr} = 1 - e^{-k_{i,corr}t}$$

$$FO = 1 - \prod_{a \in RLF} \left(1 - p_{a,corr}\right) = 1 - \exp\left[-\left(\sum_{a \in RLF} k_{a,corr}\right)t\right]$$

where $RLF$ denotes the set of band indices corresponding to the thymines labeled $T_1$ through $T_9$ in **Fig. 2B**, combining data from both the NTS-radiolabeled and TS-radiolabeled experiments. Note that the PRI metric is subject to increased uncertainty as $p_i$ approaches 1, where the slope of $\ln(\frac{1}{1-p_i})$ approaches infinity.
